# Microglial senescence contributes to female-biased neuroinflammation in the aging mouse hippocampus: implications for Alzheimer’s disease

**DOI:** 10.1101/2023.03.07.531562

**Authors:** Sarah R. Ocañas, Kevin D. Pham, Jillian E.J. Cox, Alex W. Keck, Sunghwan Ko, Felix A. Ampadu, Hunter L. Porter, Victor A. Ansere, Adam Kulpa, Collyn M. Kellogg, Adeline H. Machalinski, Ana J. Chucair-Elliott, Willard M. Freeman

**Affiliations:** Genes & Human Disease Program, Oklahoma Medical Research Foundation, Oklahoma City, OK USA; Department of Physiology, University of Oklahoma Health Sciences Center, Oklahoma City, OK USA; Graduate Program in Biomedical Sciences, University of Oklahoma Health Sciences Center, Oklahoma City, OK; Department of Biochemistry and Molecular Biology, University of Oklahoma Health Sciences Center, Oklahoma City, OK USA; Oklahoma City Veterans Affairs Medical Center, Oklahoma City, OK USA

**Author notes:** **To whom correspondence should be addressed:** Willard M. Freeman, Genes & Human Disease Program, Oklahoma Medical Research Foundation, 825 NE 13^th^ Street, Oklahoma City, OK 73104, USA. **E-mail address:**.

**Keywords:** Microglia, sex effects, sex divergence, senescence, neuroinflammation, hippocampus, brain aging, transcriptomics, disease-associated microglia, Alzheimer’s disease

## Abstract

**Background:** Microglia, the brain’s principal immune cells, have been implicated in the pathogenesis of Alzheimer’s disease (AD), a condition shown to affect more females than males. Although sex differences in microglial function and transcriptomic programming have been described across development and in disease models of AD, no studies have comprehensively identified the sex divergences that emerge in the aging mouse hippocampus. Further, existing models of AD generally develop pathology (amyloid plaques and tau tangles) early in life and fail to recapitulate the aged brain environment associated with late-onset AD. Here, we examined and compared transcriptomic and translatomic sex effects in young and old murine hippocampal microglia.

**Methods:** Hippocampal tissue from C57BL6/N and microglial NuTRAP mice of both sexes were collected at young (5-6 month-old [mo]) and old (22-25 mo) ages. Cell sorting and affinity purification techniques were used to isolate the microglial transcriptome and translatome for RNA-sequencing and differential expression analyses. Flow cytometry, qPCR, and imaging approaches were used to confirm the transcriptomic and translatomic findings.

**Results:** There were marginal sex differences identified in the young hippocampal microglia, with most differentially expressed genes (DEGs) restricted to the sex chromosomes. Both sex chromosomally-and autosomally-encoded sex differences emerged with aging. These sex DEGs identified at old age were primarily female-biased and enriched in senescent and disease-associated microglial signatures. Normalized gene expression values can be accessed through a searchable web interface (https://neuroepigenomics.omrf.org/). Pathway analyses identified upstream regulators induced to a greater extent in females than in males, including inflammatory mediators IFNG, TNF, and IL1B, as well as AD-risk genes TREM2 and APP.

**Conclusions:** These data suggest that female microglia adopt disease-associated and senescent phenotypes in the aging mouse hippocampus, even in the absence of disease pathology, to a greater extent than males. This sexually divergent microglial phenotype may explain the difference in susceptibility and disease progression in the case of AD pathology. Future studies will need to explore sex differences in microglial heterogeneity in response to AD pathology and determine how sex-specific regulators (i.e., sex chromosomal or hormonal) elicit these sex effects.

## Background

There are profound sex differences in immune responses across the lifespan, with females displaying more robust innate and adaptive immune responses. While strong immunity in females protects against infectious diseases and likely plays a role in reproductive fitness, females also have a higher incidence of autoimmunity and “inflammaging” (chronic inflammation with aging) (1, 2). Sex differences in neuroimmunity have been implicated in sex biases observed in age-related neurodegenerative diseases (i.e., Alzheimer’s disease (AD)), suggesting a potential role for microglia in driving sexually divergent effects. In addition, genome-wide association studies (GWAS) (7-10) have identified AD risk variants within genes involved in innate immunity (i.e., SPI1, TREM2) (11)), shifting the field to examine a potential causative role of microglia in the pathogenesis of AD.

Microglia are yolk-sac-derived macrophages that colonize the brain during early embryonic development. As the only resident immune cells in the brain parenchyma, microglia serve diverse roles across the lifespan, including: synaptic pruning during development, surveillance for pathogens, and clearance of debris (i.e., apoptotic cells), among others. Single-cell analyses of microglia from young mouse brains (E14.5, P4/5. P100) failed to identify major sex differences in microglial subpopulations (3). These findings align with results from our group, which suggest that sex differences in the neuroinflammatory signaling pathways emerge with age in the mouse hippocampus, and that sex differences in youth are quite limited outside sex-chromosomal encoded dimorphisms (i.e., *Xist*) (4, 5). However, to our knowledge, no studies have examined the transcriptomic and translatomic signatures of sex differences specifically in the microglia of the young and aged mouse hippocampus.

In the aging brain, microglia adopt heterogeneous functional states defined by their motility, phagocytic capacity, transcriptional programming, cell surface marker expression, and secretory phenotype. Ultimately, microglial proliferation can lead to replicative senescence, which has a phenotype that mirrors the disease-associated state observed in mouse models of AD (6). Senescent cell characteristics include: 1) cell cycle arrest, 2) resistance to apoptosis, and 3) upregulation of the senescence-associated secretory phenotype (SASP) factors. We hypothesize that female microglia have a greater burden of inflammation and senescence compared to males, thus contributing to the sex biases observed in age-related brain disease.

To test our hypothesis, we isolated and analyzed hippocampal microglial transcripts from young and old mice of both sexes using two approaches validated to minimize *ex vivo* activational artifacts (7, 8). We found that female microglia upregulate neuroinflammatory signaling pathways in the aging brain more than males. Female microglia also display a robust senescent and disease-associated phenotype with aging which is less evident in male microglia. These data suggest that a sex-specific stimulus (i.e., sex hormones or sex chromosomes) interact with aging to drive female microglia to a greater senescent phenotype than males, contributing to the heightened pro-inflammatory environment in the aged female brain. This sexually divergent microglial reactivity may contribute to the female-biased prevalence and progression of age-related brain diseases, such as AD. To provide a tool for assessing microglial sex differences to the field, normalized gene expression levels for the transcriptomic and translatomic analyses have been deployed to a searchable web interface (https://neuroepigenomics.omrf.org/).

## Materials and Methods

### Experimental animals

All animal procedures were approved by the Institutional Animal Care and Use Committee at the Oklahoma Medical Research Foundation and performed in accordance with the National Institutes of Health (NIH) Guide for the Care and Use of Laboratory Animals. All mice were housed at the Oklahoma Medical Research Foundation under SPF conditions in a HEPA barrier environment on a 14/10 light/dark cycle (lights on at 6:00 am). Male and female C57BL/6N mice were obtained from the NIA aging colony at 3 and 22 months of age (mo) and aged to 6 and 25 mo, respectively (n=3-5/sex/age). Cx3cr1-cre/ERT2^+/+^ males (Stock #020940) (9) were mated with NuTRAP^flox/flox^ females (Stock #029899) (10) to generate the desired progeny, Cx3cr1-cre/ERT2^+/wt^; NuTRAP^flox/wt^ (Cx3cr1-NuTRAP), as previously performed (7). The NuTRAP allele labels microglial ribosomes with eGFP and nuclei with biotin/mCherry, in a Cre-dependent manner (10). Cre-negative controls (Cx3cr1-cre/ERT2^wt/wt^ NuTRAP^flox/wt^) were used in flow cytometric and immunohistochemical experiments. Male and female Cx3cr1-NuTRAP mice were aged to 6 and 22 mo before tissue collection (n=5-7/sex/age) for RNA-Seq. A second cohort of male and female Cx3cr1-NuTRAP mice were aged to 8-13 mo and 23-26 mo (n=4/sex/age) for TRAP-RT-qPCR and INTACT telomere assays. DNA was extracted from mouse ear punch samples for genotyping. Genotyping was performed using standard PCR detection of Cx3cr1-cre/ERT2 (Jax protocol 27232; primers: 20669, 21058, 21059) and NuTRAP floxed allele (Jax protocol 21509; primers: 21306, 24493, 32625, 32626), as previously described (7). Mice were euthanized by cervical dislocation, followed by rapid decapitation, in line with the AVMA Guidelines for the Euthanasia of Animals.

### Tamoxifen (TAM) induction of cre recombination in Cx3cr1-NuTRAP mice

At 3-5 mo, mice received a daily intraperitoneal injection of TAM solubilized in 100% sunflower seed oil by sonication (100 mg/kg body weight, 20 mg/ml stock solution, #T5648; Millipore Sigma) for five consecutive days, as previously performed (7, 8, 11). Brain tissue was harvested at least 3 months following TAM administration to “wash-out” labeling of circulating monocytes (8).

### Preparation of Single-Cell Suspension from Mouse Brain

Brain tissue was isolated from C57BL6/N or Cx3cr1-NuTRAP mice. The hippocampus or cortex was rinsed in D-PBS, minced, and placed into gentleMACS C-tubes (#130-093-237, Miltenyi Biotec), and processed to create a single-cell suspension using the Adult Brain Dissociation Kit (#130-107-677, Miltenyi Biotec) supplemented with Actinomycin D (#A1410, Sigma-Aldrich), Triptolide (#T3652, Sigma-Aldrich), and Anisomycin (#A9789, Sigma-Aldrich) and homogenized on the gentleMACS Octo Dissociator System (#130-095-937, Miltenyi Biotec), as previously described (12). Following debris removal (#130-109-398, Miltenyi Biotec), cells were resuspended in 1 mL 0.5% BSA in D-PBS (#130-091-376, Miltenyi Biotec) and filtered through a 35 µm filter (#352235, Fisher Scientific).

### CD11b Magnetic Labeling and Separation (CD11b-MACS)

Hippocampal cells from C57BL6/N mice were pelleted at 300 x g for 10 minutes at 4°C then resuspended in 90 µL of 0.5% BSA in D-PBS and 10 µL of CD11b (Microglia) MicroBeads (#130- 093-634, Miltenyi Biotec) followed by thorough mixing. The cells were then labeled by incubating with the CD11b-beads for 15 minutes at 2-8°C. After incubation, labeled cells were washed in 1 mL of 0.5% BSA in D-PBS and then centrifuged at 300 x g for 10 minutes. Cells were resuspended in 700 µL of 0.5% BSA in D-PBS. 200 µL of the cell suspension was reserved as the CD11b-MACS Input for flow cytometric and RNA-Seq analyses. The remaining 500 µL was processed on the autoMACS Pro Separator (#130-092-545, Miltenyi Biotec). After priming the autoMACS instrument, the sample and collection tubes were placed in a cold Chill5 tube rack (#130-092-951, Miltenyi Biotec), collecting both positive and negative fractions. The double-positive selection (Posseld) program was used to elute highly pure CD11b^+^ cells. 50 µL of the positive fractions were reserved for flow cytometric analysis. The remaining cells were pelleted at 1000 x g for 10 minutes at 4°C and lysed in 350 µL of Buffer RLT (Qiagen) supplemented with 3.5 µL of 2-β mercaptoethanol (#444203, Sigma) before being stored at −80°C.

### Antibody Labeling for Flow Cytometry

Reserved cells from the CD11b-MACS Input were split into four 50 µL aliquots for antibody labeling. The first aliquot was untreated as an “Unstained” control; the second aliquot containing 1 µL of CD11b-APC antibody (#130-113-793, Miltenyi Biotec); the third aliquot containing 1 µL of CD45-VioBlue antibody (#130-110-802, Miltenyi Biotec); and the fourth aliquot containing 1 µL each of CD11b-APC antibody and CD45-VioBlue antibody. The reserved cells from the positive fraction were also stained with 1 µL each of CD11b-APC antibody and CD45-VioBlue antibody. All samples were mixed well and incubated for 10 minutes in the refrigerator (2-8°C) protected from light. Following incubation, cells were washed with 1 mL of 0.5% BSA in D-PBS and pelleted at 300 x g for 10 minutes, and then resuspended in 250 µL of 0.5% BSA in D-PBS for flow cytometric analyses.

### Flow Cytometry Analysis of CD11b-MACS purity

Analyses of the cells labeled in the previous section were conducted on a MACSQuant Analyzer 10 Flow Cytometer (#130-096-343, Miltenyi Biotec). Post-sort purity was assessed by the percent of CD11b^+^CD45^+^ singlets using MACSQuantify v2.13.0 software. Following analysis, all four aliquots from the CD11b-MACS Input were combined, pelleted at 1000 x g for 10 minutes at 4°C and lysed with 350 µL of Buffer RLT supplemented with 3.5 µL of 2-β mercaptoethanol before storing at −80°C.

### TRAP isolation from Cx3cr1-NuTRAP hippocampus (Cx3cr1-TRAP)

Hippocampal tissue was isolated from Cx3cr1-NuTRAP mice of both sexes at 6 and 22 mo (n=5- 7/sex/age). The hippocampi from both hemispheres were minced into small pieces and homogenized in 100 µL ice-cold TRAP homogenization buffer (50 mM Tris, pH 7.4; 12 mM MgCl_2_; 100 mM KCl; 1% NP-40; 1 mg/ml sodium heparin; 1 mM DTT; 100 μg/ml cycloheximide [#C4859-1ML, Millipore Sigma]; 200 units/ml RNaseOUT Recombinant Ribonuclease Inhibitor [#10777019; Thermo Fisher Scientific]; 1× complete EDTA-free Protease Inhibitor Cocktail [#11836170001; Millipore Sigma] with a Kimble pellet pestle mixer [#749540-0000; DWK Life Sciences), as previously described (7, 8). Volume was brought up to 1.5 mL with TRAP homogenization buffer. Homogenates were centrifuged at 12,000 × *g* for 10 min at 4°C. After centrifugation, 100 μL of the cleared supernatant was saved as TRAP Input. The remaining supernatant was incubated with 5 μg/μL of anti-GFP antibody (ab290; Abcam) at 4°C with end-over-end rotation for 1 hour. A volume of 50 µL per sample of Protein G Dynabeads (#10003D; Thermo Fisher Scientific) was washed three times in 1-mL ice-cold low-salt wash buffer (50 mM Tris, pH 7.5; 12 mM MgCl_2_; 100 mM KCl; 1% NP-40; 100 μg/ml cycloheximide; 1 mM DTT) and then resuspended in the original volume of low salt wash buffer. 50 µL of washed beads were added to each cleared homogenate and incubated at 4°C with end-over-end rotation overnight. Following overnight incubation, the beads and GFP-bound polysomes were washed three times with 0.5 mL of high-salt wash buffer (50 mM Tris, pH 7.5; 12 mM MgCl_2_; 300 mM KCl; 1% NP-40; 100 μg/mL cycloheximide; 2 mM DTT) using a magnetic stand for separation. After the last wash, 350 μL of Buffer RLT (Qiagen) was supplemented with 3.5 μl of 2-β mercaptoethanol (#444203, Millipore Sigma) and added to the washed beads. Samples were incubated with mixing on a ThermoMixer (Eppendorf) for 10 minutes at room temperature. The beads were magnetically separated, and the supernatant (Positive Fraction) was transferred to a new tube. RNA was isolated using the RNeasy Mini kit (#74104, QIAGEN) according to the manufacturer’s instructions. RNA was quantified with a Nanodrop One^c^ spectrophotometer (#ND-ONEC-W, Thermo Fisher Scientific), and its quality was assessed by HSRNA ScreenTape (#5067-5579, Agilent Technologies) with a 4150 TapeStation analyzer (#G2992AA, Agilent Technologies).

### RNA Extraction

Input and positive fraction samples from CD11b-MACS and Cx3cr1-TRAP were removed from storage and thawed on ice. The lysate was then loaded onto a QIAshredder spin column (#79656, Qiagen) and placed in a 2 mL collection tube before being centrifuged for 2 minutes at full speed. RNA was then isolated using an AllPrep Mini Kit (#80204, Qiagen) according to the manufacturer’s instructions. RNA was quantified with a Nanodrop One^C^ spectrophotometer (#ND-ONEC-W, ThermoFisher Scientific) and quality-assessed by HSRNA ScreenTape (#5067-5579, Agilent Technologies) with a 4150 TapeStation analyzer (#G2992AA, Agilent Technologies).

### Library Construction and RNA Sequencing (RNA-seq)

Directional RNA-Seq libraries were made according to the manufacturer’s protocol from 5-100 ng RNA. Poly-adenylated RNA was captured using NEBNext Poly(A) mRNA Magnetic Isolation Module (#NEBE7490L, New England Biolabs) and then processed using NEBNext Ultra II Directional Library Prep Kit for Illumina (#NEBE7760L, New England Biolabs) for the creation of cDNA libraries. Library sizing was performed with HSD1000 ScreenTape (#5067-5584, Agilent Technologies) and quantified by Qubit dsDNA HS Assay Kit (#Q32851, ThermoFisher Scientific) on a Qubit 4 Fluorometer (#Q33226, ThermoFisher Scientific). The libraries for each sample were pooled at 4 nM concentration and sequenced using an Illumina NovaSeq 6000 system (SP PE50bp, S4 PE150).

### RNA-Seq Analysis

After sequencing, reads were aligned against the mm10 transcriptome and genome (UCSC) in StrandNGS software (v4.0; Strand Life Sciences). Parameters for alignment and filtering included: adapter trimming, fixed 2-bp trim from 5′ and 2 bp from 3′ ends, a maximum number of one novel splice allowed per read, a minimum of 90% identity with the reference sequence, a maximum 5% gap, and trimming of 3′ ends with Q < 30. Alignment was performed directionally with Read 1 aligned in reverse and Read 2 in the forward orientation, as previously (8). Binary alignment map (BAM) files were exported from StrandNGS software for further analysis. The Rsubread package (13) was used to quantify the read counts using the featureCount command. Normalization and differential expression calling was then performed using edgeR (14) with the quasi-likelihood pipeline (15). Differentially expressed genes (DEGs) (GLM QLF<0.1, |FC|>1.25) were identified for each of the comparisons (YF v YM, OF v OM, OF v YF, OM v YM) as well as the interaction term. DEGs were uploaded to Ingenuity Pathway Analysis software (IPA; Qiagen) to assess the biological significance of differential expression. Heatmaps of DEGs and pathway activation z-scores were made using Morpheus software (https://software.broadinstitute.org/morpheus). Volcano plots were made in R using the EnhancedVolcano package. Upset plots were created using the UpSetR package. Gene set enrichment analysis (GSEA) was run using GSEA v4.1.0 (16). For the differential transcript analysis of the *Cdkn2a* gene, StrandNGS software was used for quantitation.

### Assessment of Cx3cr1-NuTRAP Cre specificity by flow cytometry

Adjacent cortex tissue from the same Cx3cr1-NuTRAP mice used for the hippocampal TRAP-Seq analyses were collected for flow cytometric analyses. After creating a single-cell suspension (as described above), cells were labeled with CD11b and CD45 fluorescent antibodies. Analyses of the labeled cells were conducted on a MACSQuant Analyzer 10 Flow Cytometer (#130-096-343, Miltenyi Biotec). The proportion of eGFP^+^ cells that co-labelled with CD11b^+^CD45^+^ was assessed and compared between groups using MACSQuantify v2.13.0 software.

### RT-qPCR analysis from TRAP-isolated RNA

Relative gene expression levels were quantified by qPCR as previously described (17, 18). Starting with 20 ng of TRAP-isolated RNA, cDNA was synthesized using the ABI High-Capacity cDNA Reverse Transcription Kit (Applied Biosystems Inc., Foster City, CA). qPCR was performed with gene-specific primer probe fluorogenic exonuclease assays (PrimeTime qPCR Probe Assays, Integrated DNA Technologies, Coralville, Iowa, **Supplemental Table 1**) and the QuantStudio 5 Real-Time PCR System (Applied Biosystems). Relative gene expression (RQ) was calculated with Expression Suite v 1.3 software using the 2^−ΔΔ^Ct analysis method with *Hprt* (Mm01324427_m1, TaqMan, Life Technologies, Waltham, MA) as an endogenous control. Statistical analysis of the qPCR data was performed using GraphPad Prism 9 (San Diego, CA). Two-way ANOVA was performed followed by the Tukey’s multiple comparison test (p< 0.05).

### Creation of nuclear suspension

The hippocampi from both hemispheres of Cx3cr1-NuTRAP mice were cut into four pieces and added to the pre-cooled gentleMACS C-tube (#130-096-334, Miltenyi Biotec) pre-filled with 2 mL ice-cold Nuclei Extraction Buffer (#130-128-024, Miltenyi Biotec) supplemented with 0.2 U/μL RNase inhibitor (#2489605, Thermo Fisher Scientific) (referred to as Nuclei Extraction Buffer hereafter). The gentleMACS C-tube was mounted onto the gentleMACS Dissociator (#130-095- 937, Miltenyi Biotec) and ran gentleMACS Program 4C_nuclei1. After termination of the program, the nuclear suspension was filtered through the MACS SmartStrainer (100 μm) (#130-098-463, Miltenyi Biotec) into a 15 mL conical tube. The filter was washed with 2 mL ice-cold Nuclei Extraction Buffer for maximum nuclei recovery. The filter was discarded, and the nuclear suspension was centrifuged at 300 x g at 4˚C for 5 minutes. The supernatant was aspirated, and the nuclei pellet was resuspended in 2 mL of ice-cold resuspension buffer (1 M HEPES [#J16924- AE, ThermoFisher Scientific]; 5 M NaCl [#01094764, ThermoFisher Scientific]; 90 mM KCl [#01276444, ThermoFisher Scientific]; 2 mM EDTA [#01216827, ThermoFisher Scientific]; 0.5 mM EGTA [#40520008-1, bioWORLD]) with 1X protease inhibitor (#1861278, ThermoFisher Scientific) (referred to as NPB hereafter) by slowly and gently pipetting the sample up and down for ten times. The nuclear suspension was filtered through the MACS SmartStrainer (30 μm) (#130-098-458, Miltenyi Biotec) into a 15 mL conical tube. The sample was centrifuged at 500 x g for 5 minutes at 4˚C. The supernatant was then aspirated, and the nuclear pellet was resuspended with 200 μL Nuclei EZ Storage buffer (#NUC-101, Millipore Sigma) by briefly vortexing. Twenty microliters of the sample were reserved as nuclei input sample. The remaining nuclei sample was filled back to 2 mL volume with ice-cold NPB with 1X protease inhibitor and subjected to the INTACT protocol.

### INTACT-isolation of hippocampal microglial DNA

INTACT isolation of microglial nuclei from Cx3cr1-NuTRAP hippocampi was performed as previously described, with all steps carried out at 4˚C (7). Briefly, each hippocampal nuclei suspension was subjected to a 40-minute incubation with 30 µL pre-washed M-280 Streptavidin Dynabeads (#11205, ThermoFisher Scientific). The mixture of nuclei and beads in each sample was placed on a Dyna-2 magnetic stand (#12321.D, Thermofisher Scientific) for 3 minutes, to separate biotinylated nuclei, corresponding to microglia (positive fraction), from nuclei from other cell types (negative fraction). After removal of the supernatant (negative fraction), the positive nuclear fraction was washed in 1 mL of NPB three times, and resuspended in 30 μL of NPB. Both input and positive fractions were stored at −80˚C until the DNA extraction was performed.

### gDNA Extraction

Input and positive fraction nuclear samples from the Cx3cr1-NuTRAP mice were removed from storage and thawed on ice. DNA was then isolated using AllPrep DNA/RNA Mini Kit (#80204, Qiagen) according to the manufacturer’s instructions. DNA was quantified with a Nanodrop One^C^ spectrophotometer (#ND-ONEC-W, ThermoFisher Scientific), Qubit4 Fluorometer (#2322619089537, ThermoFisher Scientific), and quality-assessed by Genomic DNA ScreenTape (#5067-5365, Agilent Technologies) with a 4150 TapeStation analyzer (#G2992AA, Agilent Technologies).

### Telomere length assay

INTACT-isolated DNA from the input and positive fractions were normalized to 0.5 ng/µL. Relative telomere length was measured using a previously described RT-qPCR method with minor modifications (19). Briefly, 1 ng DNA from each sample was combined with PowerUp SYBR Green Master Mix (#A25741, Thermofisher) and specific primers for either 36B4 (control) or telomere sequences (**Supplemental Table 1**). Samples were run in triplicate on the Quantstudio5 using the following cycling conditions: denaturation at 95°C for 10 minutes followed by 40 cycles of 95°C for 15 seconds and 60°C for 1 minute. Telomere length data were normalized to the 36B4 control using the 2^-ΔΔCt^ method.

### Immunohistochemistry

For immunohistochemistry (IHC), mouse brains were harvested from both female and male Cx3cr1-NuTRAP mice and hemisected. Samples were fixed in 4% PFA for 4 hours and cryoprotected in PBS containing 30% sucrose. Samples were then cryo-embedded in Optimal Cutting Temperature medium (Tissue-Tek, #4583) and sectioned at a thickness of 12- micrometers using the Cryostar NX70 (ThermoFisher Scientific) (7, 8). For analysis of CD11b and CD11c expression, cryo-sectioned samples were thawed at room temperature for 5 min and mounted onto a Shandon Sequenza Immunostaining Center Slide Rack (#73310017, Epredia, Kalamazoo, MI) for immunotaining. The slides were then washed three times with 1% Triton x- 100 in PBS to rehydrate, followed by blocking with 10% normal donkey serum for 1 hour at room temperature. Primary antibodies (**Supplemental Table 1**) were added to an incubation buffer (1% BSA, 1% Normal Serum, 0.3% Triton X-100 in PBS) according to the dilutions indicated below and incubated for 22 hours at 4°C. On the following day, slides were washed three times with 1x PBS, followed by a 1-hour incubation with secondary antibodies (**Supplemental Table 1**). Slides were washed three times before a 5-minute incubation with DAPI followed by one last wash with 1x PBS before mounting.

A Zeiss LSM 710 Confocal microscope at the OMRF Imaging Core was used to capture the dentate gyrus region of sagittal brain sections. The images were collected at 20X magnification using the Z-stack (1.00 µm step size and a total of 15-17 slices) and tile scan feature and stitched in their native format (.czi). Images were processed using ImageJ (Fiji) and then exported as .png files for figure assembly. The instrument settings used for raw capture and image processing are disclosed in **Supplemental Table 1**. Separated channels for the images presented in **Figure 6C-D** are given as **Supplemental Figure 1**.

### Statistics

Boxplots were generated using GraphPad Prism software. The box represents the median +/- interquartile range with whiskers extending to the minimum and maximum values within the dataset. Two-way ANOVA’s were performed with factors of age (O v Y) and sex (F v M) and the interaction of age:sex. Post-hoc comparisons were made and corrected for using either Tukey’s or Sidak’s multiple comparisons tests, as indicated in the figure legends.

## Results

This study aimed to analyze the sex-differential gene expression response of female and male hippocampal microglia to brain aging. To this end, mice of both sexes were collected at young (5- 6 months old [mo]) and old (22-25 mo) ages (**Figure 1A**). To ensure rigor and reproducibility, microglial RNA was enriched from acutely isolated hippocampal tissue by two previously validated methods to minimize *ex vivo* activational confounds of microglial isolation (8): 1) CD11b^+^ magnetic-activated cell sorting (CD11b-MACS) from C57BL/6N mice with the inclusion of transcription and translation inhibitors during cell preparation, and 2) Translating Ribosome Affinity Purification from Cx3cr1-NuTRAP mice (7) (Cx3cr1-TRAP) which allow for transcriptome and translatome assessments, respectively.

**Figure 1.**
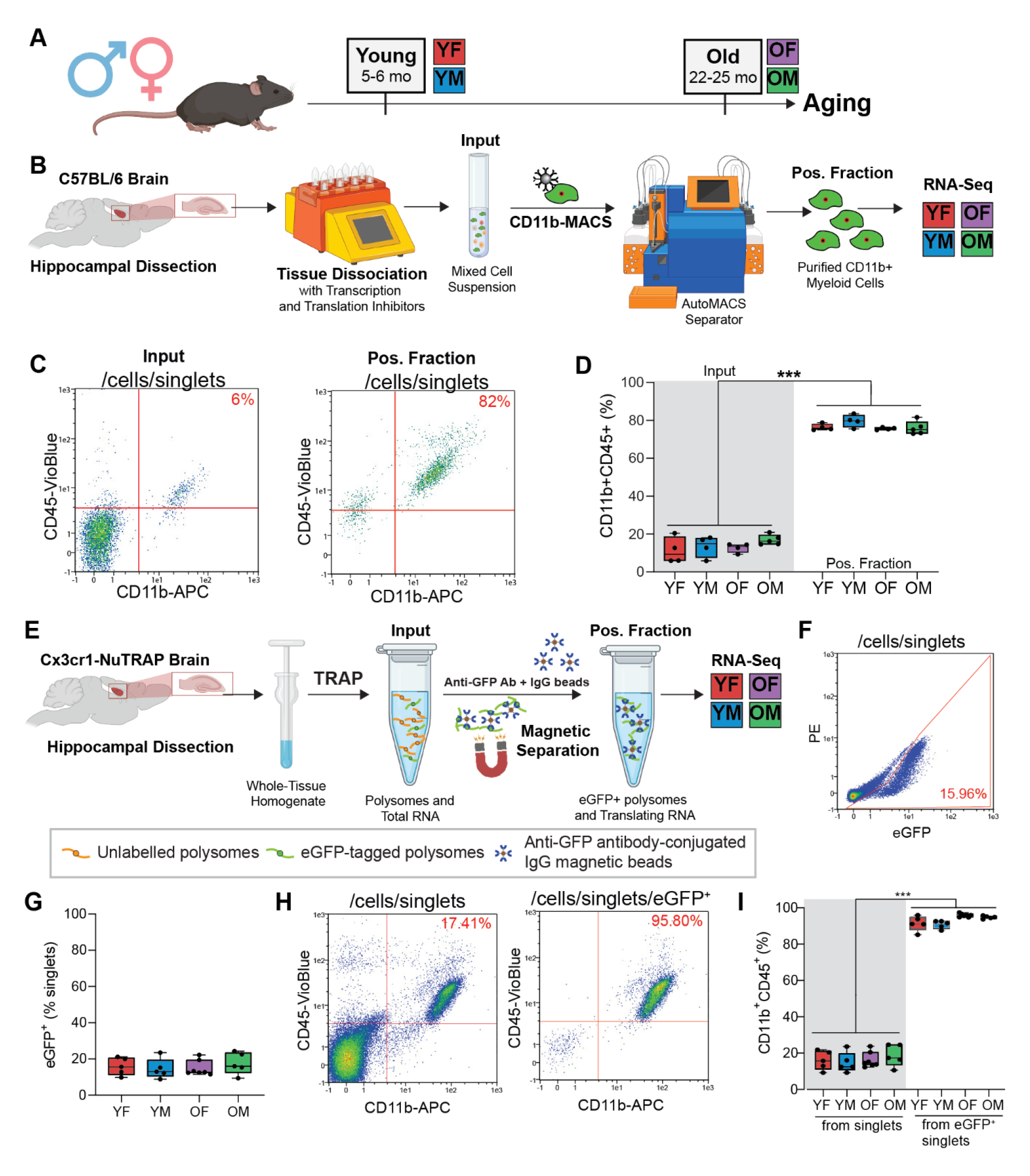
Isolation of hippocampal microglial transcripts by CD11b-MACS and Cx3cr1-TRAP. **A)** Schematic of experimental grouping. Male and female C57BL/6N or Cx3cr1-NuTRAP mice were collected at young (5-6 months) and old (22-25 months) timepoints to form four groups: 1) young female (YF), 2) young male (YM), 3) old female (OF), and 4) old male (OM). **B)** Isolation of hippocampal microglia by CD11b-MACS. The hippocampus from YF, YM, OF, and OM C57BL6/N mice was dissociated by enzymatic and mechanical dissociation with transcription and translation inhibitors. The single-cell suspension was labeled with CD11b microbeads prior to magnetic separation. The CD11b^+^ fraction was then analyzed by flow cytometry. **C)** Representative flow cytometry plots of the CD11b and CD45 immunoreactivity from the CD11b-MACS input and positive (pos.) fractions. **D)** Quantitation of the percentage of singlets that were CD11b^+^CD45^+^ from the CD11b-MACS input and pos. fraction (Two-way ANOVA, main effect MACS fraction [Input v. Pos. Fraction], ***p<0.001). **E)** Isolation of hippocampal microglial translatome by Cx3cr1-TRAP. Mouse hippocampus was homogenized in TRAP lysis buffer containing translation inhibitors. eGFP-labeled polysomes and associated translating RNA (from Cx3cr1^+^ cells) were magnetically separated using an eGFP antibody and IgG beads. RNA from the pos. fraction was used to generate stranded RNA-Seq libraries for assessment of the microglial translatome. **F-I)** Cx3cr1-NuTRAP cortex samples were used to assess the cell specificity of cre-mediated induction of the NuTRAP allele by flow cytometry. **F)** Representative flow cytometry plot of eGFP^+^ singlets from Cx3cr1-NuTRAP cortex. **G)** Quantitation of the percent eGFP^+^ singlets from young and old Cx3cr1-NuTRAP cortex from both sexes. **H)** Representative flow cytometry plots of the percent CD11b^+^CD45^+^ singlets (left) and eGFP^+^ singlets (right). **I)** Quantitation of the percent CD11b^+^CD45^+^ singlets (left) and eGFP^+^ singlets (right).

### CD11b-MACS of C57BL6/N mouse hippocampus results in highly pure microglia for transcriptomic analyses

At young (5-6 mo) and old (22-25 mo) ages, C57BL6/N mice of both sexes were collected for CD11b-MACS isolation from acutely isolated hippocampus. Following hippocampal dissection, the tissue was enzymatically and mechanically dissociated with the inclusion of transcription and translation inhibitors to prevent *ex vivo* activational artifacts, as previously performed (8). The mixed cell suspension was then incubated with CD11b magnetic beads and magnetically separated. RNA was isolated from an aliquot of the mixed cell suspension (Input) and sorted CD11b^+^ cells (Positive Fraction), and processed for RNA-Seq analysis (**Figure 1B**). The purity of the CD11b-MACS positive fraction was assessed by flow cytometry in comparison to the input (**Figure 1C-D**). The CD11b-MACS positive fraction was composed of ∼80% CD11b^+^CD45^+^ cells, regardless of age or sex, as compared to 10-20% in the input (**Figure 1D**, Two-way ANOVA, main effect input v. positive fraction, ***p<0.001).

### Cx3cr1-TRAP captured translating microglial RNA from Cx3cr1-NuTRAP mouse hippocampus

At young (5-6 mo) and old (22-25 mo) age, Cx3cr1-NuTRAP mice of both sexes were collected for isolation of translating microglial RNA from acutely isolated hippocampus. Following hippocampal dissection, tissue was homogenized in a buffer containing translation inhibitors to maintain interaction between the translating RNA and polysomes. Separation of microglial eGFP-tagged polysomes (and associated RNA) was then conducted using the TRAP procedure (**Figure 1E**). Flow cytometry from adjacent cortex samples was conducted to confirm the identity of eGFP-labelled cells as microglia (**Figure 1F-I**). There was a distinct eGFP^+^ population across all groups, accounting for ∼15% of the total cell population (**Figure 1F-G**). Among the eGFP^+^ cells, approximately 95% co-expressed CD11b^+^CD45^+^, thereby confirming the microglial identity of the eGFP^+^ population (**Figure 1H-I**).

A previous report from our group confirmed that the CD11b-MACS and Cx3cr1-TRAP isolation techniques resulted in high levels of microglial enrichment, but may also isolate transcripts from other brain-resident macrophages (8). For ease of explanation, we will refer to these cells as microglia, however, it is important to note that there may also be contributions from non-microglial macrophages as well.

### Hippocampal microglia showed a consistent aging signature across both methods

Following the assessment of purity, microglial transcriptomic (CD11b-MACS) and translatomic (Cx3cr1-TRAP) analyses were conducted. A multidimensional scaling (MDS) plot of the CD11b-MACS transcriptomic analysis showed separation in the first component by age (**Figure 2A**). Although there was no separation of male and female microglia in the young samples, there were distinct clusters for old females (OF) and old males (OM) (**Figure 2A**), with OF being further to the right on the first component, that was associated with age, than OM. The CD11b-MACS transcriptomic analysis identified 3734 genes differentially expressed by age in females and 2810 genes differentially expressed by age in males, with 2063 age effects in common between both sexes (**Figure 2B; Supplemental Table 2**) (Quasi-likelihood (QL) F-test, FDR<0.1). For the Cx3cr1-TRAP translatomic analyses, the MDS plot showed separation in the first component by both age and sex, with young samples clustering to the left, old samples to the right, and separation by sex at both ages **(Figure 2C**). The Cx3cr1-TRAP translatomic analysis identified 1195 genes differentially expressed by age in females and 3414 genes differentially expressed by age in males, with 762 age effects in common between both sexes (**Figure 2D; Supplemental Table 3**) (QLF-test, FDR<0.1). Volcano plots of differentially expressed genes (DEGs) by age showed similar distributions across each age and method, with an over-representation of genes being upregulated with age (**Figure 2E-H**) (QLF test, FDR<0.1, |FC|>1.25). Interestingly, genes associated with senescence (i.e., *Cdkn2a*, *Spp1, C3, Ccl8*) (20) and disease-associated microglia (DAMs) (i.e., *Itgax, Clec7a, Cst7*) (21) appear among the most highly induced genes with aging in at least one of the comparisons. These data show robust age effects induced in hippocampal microglia and provide evidence that senescent and disease-associated programs are altered with age.

**Figure 2.**
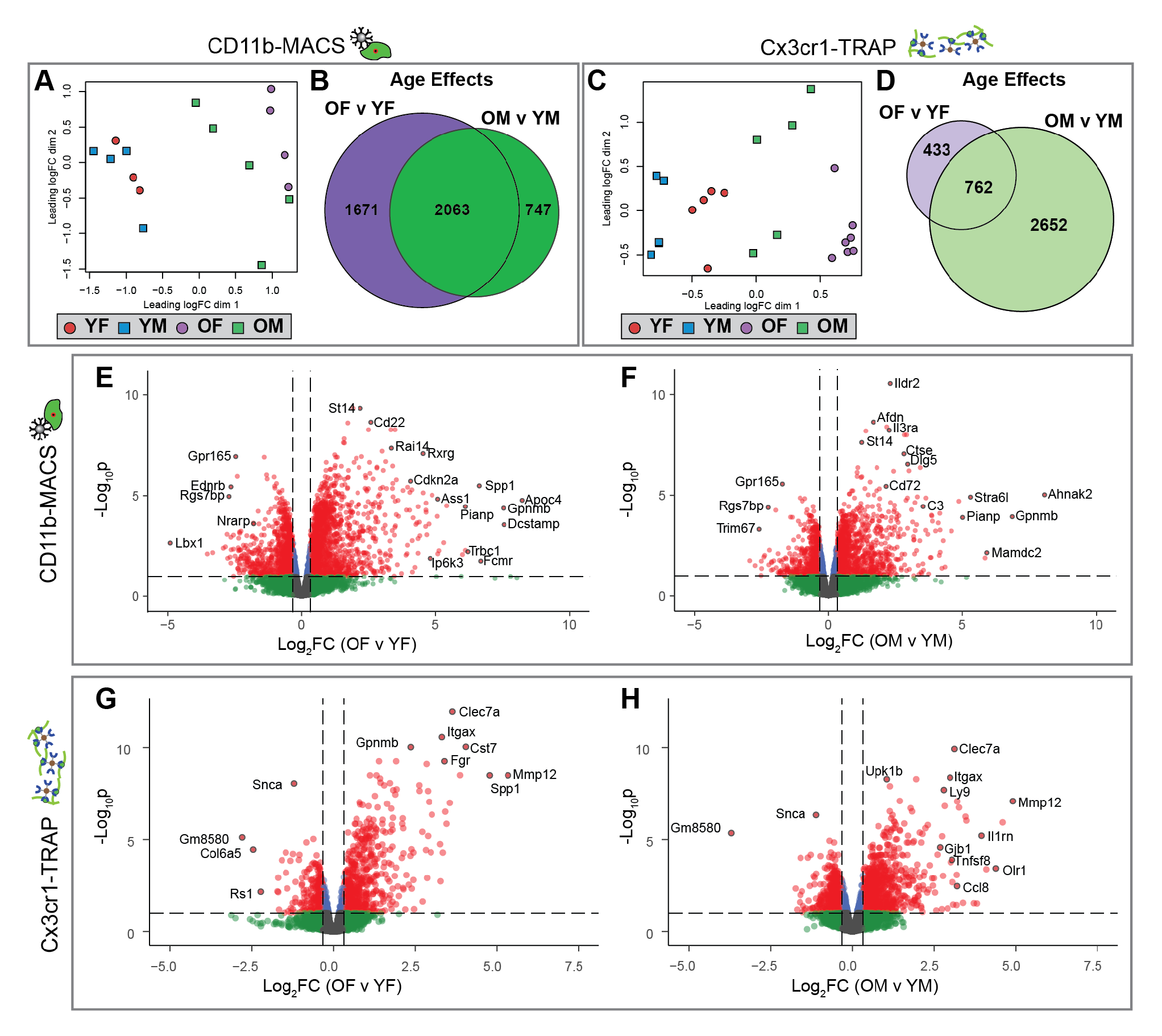
Analysis of sex-specific microglial transcriptomic (CD11b-MACS) and translatomic (Cx3cr1-TRAP) age effects from mouse hippocampus. Stranded RNA-Seq libraries were constructed from microglial RNA isolated by CD11b-MACS (n=3-5/sex/age) or Cx3cr1-TRAP (n=5-7/sex/age) for groups described in **Fig. 1A** and sequenced on an Illumina NovaSeq6000 platform. Demultiplexed fastq files were aligned and deduplicated in StrandNGS software prior to quantification (featureCounts) and differential expression calling (edgeR) in R. **A)** Multidimensional scaling (MDS) plot of CD11b-MACS gene expression shows separation in the first dimension by age. Of note, the young samples do not show separation by sex, whereas the old samples can be separated into male and female groups. **B)** Sets of differentially expressed genes (GLM QLF-test, FDR<0.1) showing an age effect in females (OF v YF) or males (OM v YM) within the CD11b-MACS dataset were compared by Venn diagram. **C)** MDS plot of Cx3cr1-TRAP gene expression shows separation in the first dimension by age and sex. **D)** Sets of differentially expressed genes (GLM QLF-test, FDR<0.1) showing an age effect in females (OF v YF) or males (OM v YM) within the Cx3cr1-TRAP dataset were compared by Venn diagram. Volcano plots of differentially expressed genes with age (GLM QLF-test, FDR<0.1, |FC|>1.25) within CD11b-MACS for **(E)** females and **(F)** males, in addition to Cx3cr1-TRAP for **(G)** females and **(H)** males.

### Sex-independent age effects in hippocampal microglial transcriptomic and translatomic analyses

To identify age effects that were common between the transcriptomic (CD11b-MACS) and translatomic (Cx3cr1-TRAP) analyses, the DEGs identified in **Figure 2E-H** were intersected (**Figure 3A; Supplemental Tables 2-3**). There were 245 age DEGs that were observed across both methods and sexes (**Figure 3A; Supplemental Table 4**). Of these 245 genes, 98% changed in the same direction with age across all comparisons, with 230 genes upregulated and 11 genes downregulated with age (**Figure 3B; Supplemental Table 4**). We identified sex-common, method-independent changes in important microglial regulators (**Figure 3C; Supplemental Table 4**). For example, pro-inflammatory mediators (i.e., TNF, IL1B, IFNG, TLR7), AD-associated genes (i.e., APP, TREM2), metabolic regulators (i.e., MTORC1, IGF1), and proliferation-associated genes (i.e., CSF1, SPI1) were upregulated with age in both sexes and across both isolation methods. Consistent with previous reports examining age-related changes in microglia (3, 22), we observed an overall increase in inflammatory signaling pathways (i.e., necroptosis, wound healing, neuroinflammation) by IPA canonical pathway analysis (**Figure 3D; Supplemental Table 4**). Additionally, analysis of disease and biological function pathways showed increased phagocytic migration and quantity of phagocytes in both sexes, which was conserved across both isolation methods (**Figure 3D; Supplemental Table 4**). Next, we examined the GO Biological Processes (hyper-geometric test, BHMTC, FDR<0.05) for genes that were consistently upregulated or downregulated across both sexes and method and also not identified as a sex effect in any comparison. The top five GO biological processes included antigen processing and presentation, lipid processing, defense response to bacterium, and response to wounding (**Figure 3E; Supplemental Table 4**). In line with previous reports from our lab (4, 23), major histocompatibility complex I (MHC-I) components *B2m*, *H2-K1*, and *H2-D1* are upregulated with age across all groups (**Figure 3F**). Within the lipid localization pathway, there is noted upregulation of apolipoprotein E (*Apoe*) and low-density lipoprotein receptor (*Ldlr*) (**Figure 3G**).

**Figure 3.**
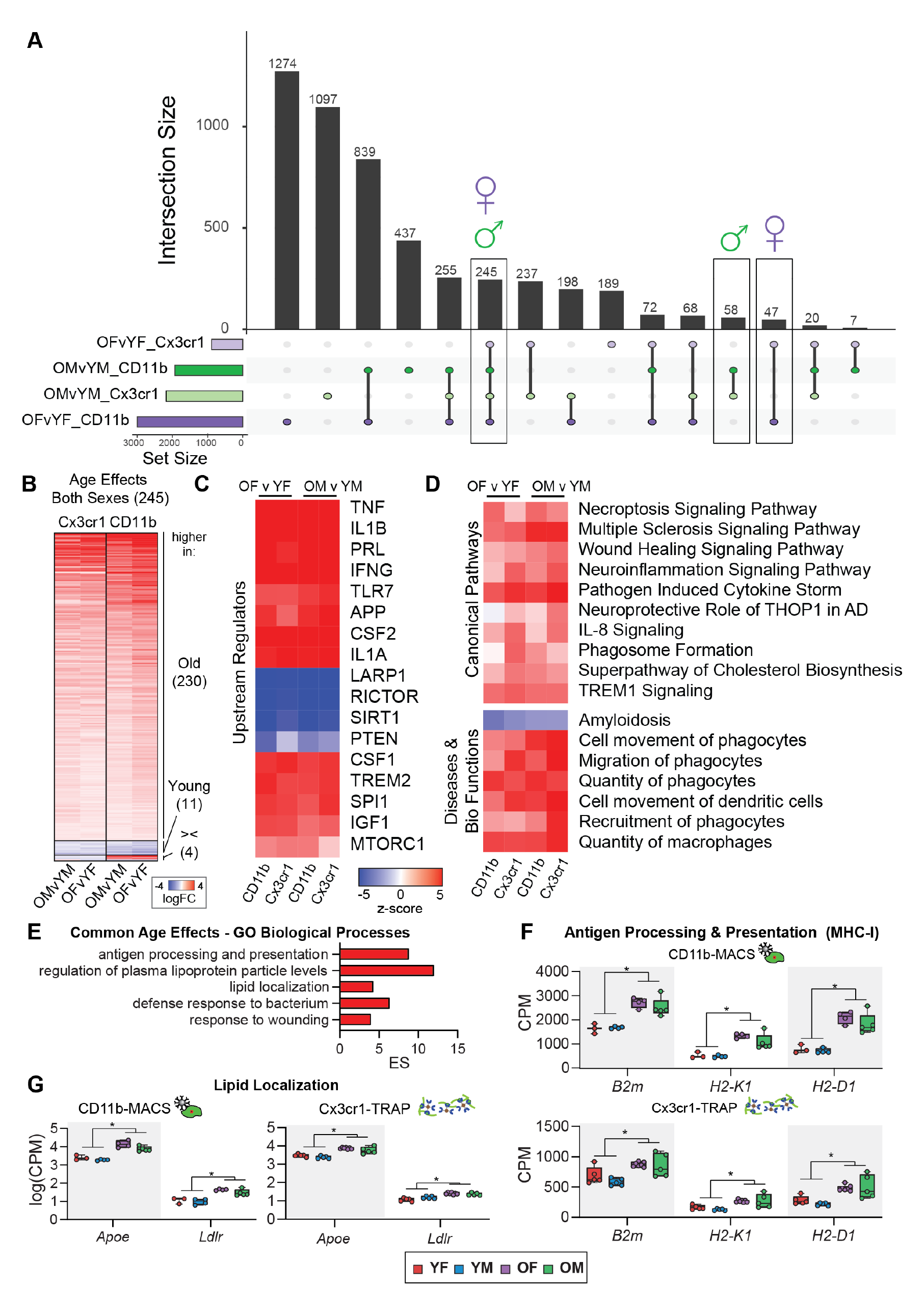
Comparison of hippocampal microglial transcriptomic age effects between CD11b-MACS and Cx3cr1-TRAP methods. **A)** Upset plot of differentially expressed genes by age with both sexes and across both methods (GLM QLF-test, FDR<0.1, |FC|>1.25). **B)** Differentially expressed genes with age that were identified in both males and females across both methods (GLM QLF-test, FDR<0.1, |FC|>1.25). **C)** Ingenuity pathway analysis identified conserved ‘Upstream Regulators’ across both sexes and methods (|z|>2, p<0.05). **D)** Ingenuity pathway analysis identified conserved ‘Canonical Pathways’ and ‘Diseases and Bio Functions’ across both sexes and methods (|z|>2, p<0.05). **E)** Top 5 GO Biological Processes for age effects altered in the same direction across both methods and sexes (Hypergeometric test, BH MTC, FDR<0.05). **F)** Gene expression (CPM) of antigen processing and presentation genes (*B2m, H2- K1, H2-D1*) (GLM QLF-test, |FC|>1.25, *FDR<0.1 for OF v YF and OM v YM comparisons). **G)** Gene expression (log_10_CPM) of lipid localization genes (*Apoe*, *Lplr*) (GLM QLF-test, |FC|>1.25, *FDR<0.1 for OF v YF and OM v YM comparisons).

The consistency of these signatures across both sexes and methods, as well as the uniformity with previously published reports provides strong confidence in the reliability of the presented methods for microglial transcriptomic and translatomic analyses.

### Sex-specific age effects in hippocampal microglial transcriptomic and translatomic analyses

There were 47 genes found to be altered only in females in both analyses (**Figure 3A**), with 24 upregulated, 21 downregulated, and 2 discordant genes across the two methods (**Figure 4A; Supplemental Table 5**). There were 58 genes identified as differentially expressed by age in only males in both analyses. Of those 58 genes, 42 genes were altered in the same direction across the transcriptomic and translatomic analyses, with 29 upregulated, 13 downregulated, and 16 discordant genes across both methods (**Figure 4B; Supplemental Table 5**). We next identified IPA upstream regulators that were differentially altered with age in only females (**Figure 4C; Supplemental Table 5**) or males (**Figure 4D; Supplemental Table 5**). Discoidin Domain Receptor 1 (DDR1) is a tyrosine kinase cell surface receptor that is activated by collagen (24) and upregulated in neurodegenerative diseases (25). In the present study, female microglia displayed upregulated DDR1 signaling with aging in both the transcriptomic and translatomic analyses (**Figure 4C; Supplemental Table 5**). DDR1 signaling was not altered in males with aging in either of the analyses. DDR1 downstream effector genes that were altered in female microglia with aging included matrix metalloproteinases (*Mmp2*, *Mmp14*), cytokines (*Ccl2*), and cellular senescence genes (*Cdkn1a*, *Cdkn2a*) (**Figure 4E**). These results suggest that female microglia may preferentially upregulate cellular senescent programs in the aged brain. Beta-galactosidase-1 (*Glb1*) and B-cell lymphoma-2 (*Bcl2*) were among the genes that were upregulated with age in females and not in males (**Figure 4A**). *Glb1* encodes for the lysosomal β-galactosidase, which is one of the most widely used markers of cellular senescence (26). *Bcl2* encodes for an apoptosis inhibitory protein and has also been associated with senescent phenotypes (27). The female-biased induction of senescent-associated genes with aging suggests that female microglia skew toward senescent phenotypes to a greater extent than males, which may account for the enhanced neuroinflammatory signatures observed in the aged female hippocampus (5). In males, ubiquitin specific protease 18 (USP18) was preferentially downregulated with age (**Figure 4D**). Downregulation of USP18 induces the expression of interferon-response genes (28), which we observe in the present study (**Figure 4H**). Interferon-induced protein with tetratricopeptide repeats 1 (*Ifit1*) and Interferon Regulatory Factor 7 (*Irf7*) were preferentially upregulated in male microglia (**Figure 4B, 4I-J**). Both *Irf7* and *Ifit1* are broadly involved in type I interferon response pathways, and are involved in antiviral innate immunity (29, 30). These results suggest that male microglia may preferentially upregulate interferon-response pathways through a USP18-dependent mechanism.

**Figure 4.**
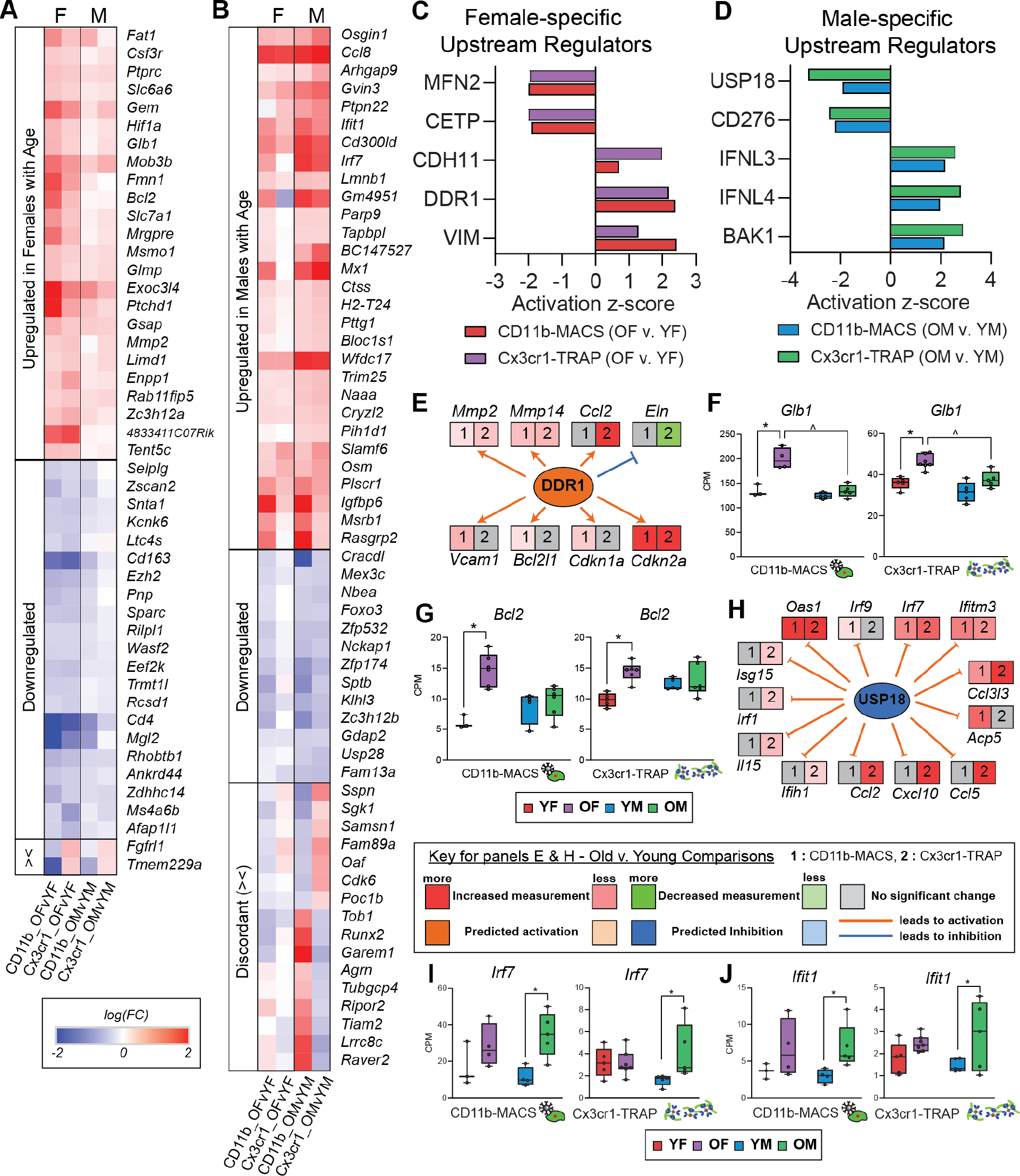
Sex-specific age effects in the microglial transcriptome and translatome. **A)** Heatmap of female-specific age effects (OF v. YF) identified as significantly upregulated or downregulated in the CD11b-MACS and Cx3cr1-TRAP analyses (GLM QLF-test, FDR<0.1, |FC|>1.25). **B)** Heatmap of male-specific age effects (OM v. YM) identified as significantly upregulated or downregulated in the CD11b-MACS and Cx3cr1-TRAP analyses (GLM QLF-test, FDR<0.1, |FC|>1.25). **C)** Select IPA upstream regulators that were identified as differentially regulated by age in females only across the CD11b-MACS and Cx3cr1-TRAP analyses (FDR<0.1). **D)** Select IPA upstream regulators that were identified as differentially regulated by age in males only across the CD11b-MACS and Cx3cr1-TRAP analyses (FDR<0.1). **E)** Expression analysis of genes downstream of IPA upstream regulator DDR1 that is altered with age only in females. **F-G)** Gene expression (CPM) of (**F**) *Glb1* and (**G**) *Bcl2* which change with age in only females (GLM QLF-test, |FC|>1.25, *FDR<0.1 age effect (O v. Y), ^FDR<0.1, sex effect (F v. M)). Boxplots represent the median +/- IQR. **H)** Expression analysis of genes downstream of IPA upstream regulator USP18 that is altered with age in males only. **I-J)** Gene expression (CPM) of **(I)** *Irf7* and **(J)** *Ifit1* which change with age in only males (GLM QLF-test, |FC|>1.25, *FDR<0.1 age effect (O v. Y), ^FDR<0.1, sex effect (F v. M)). Boxplots represent the median +/- IQR.

### Sex effects in the microglial transcriptome and translatome in the aging mouse hippocampus

We next sought to assess microglial sex effects in the mouse hippocampus within young and old mice across both transcriptomic and translatomic analyses. The CD11b-MACS transcriptomic analyses identified 8 genes that were differentially expressed by sex in both young and old mice, with 197 additional DEGs identified in the old comparison alone (**Figure 5A; Supplemental Table 2**). Of note, the eight sex DEGs at young age were all sex-chromosomally encoded. In contrast, the sex DEGs identified in the old hippocampus were a mix of autosomally-and sex-chromosomally-encoded genes. In the Cx3cr1-TRAP analyses, we identified 29 genes that were only differentially expressed in the young age group, 344 genes that were only differentially expressed in the old age group, and 23 genes that were differentially expressed across both age groups (**Figure 5B; Supplemental Table 3**). We then explored the interaction between genes that were differentially expressed by sex in the transcriptome and translatome of aging microglia (**Figure 5C; Supplemental Table 2-3**). Seven sex DEGs were identified in young and old samples from both the transcriptomic and translatomic analyses, with three being X chromosomally-encoded transcripts (*Kdm6a*, *Eif2s3x, Xist*) and four Y chromosomally-encoded (*Kdm5d, Eif2s3y, Uty, Ddx3y*), consistent with sexually dimorphic genes identified in previous studies from mouse hippocampus (17). Heatmaps of the relative gene expression (counts per million – CPM) showed high expression of the X-chromosomally encoded genes (*Kdm6a*, *Eif2s3x, Xist*) in females when compared to males (**Figure 5D-E**). *Kdm6a* encodes for the lysine-specific demethylase 6A histone modifier, which selectively demethylates H3K27, serving an important role in modulating the epigenome (31). Recently, *Kdm6a* has been shown to play an important role in hippocampal-dependent cognitive function (32-34). It stands to reason that the sexually dimorphic expression of *Kdm6a* may establish differences in the sex-specific microglial epigenomic landscape and mediate differential responses to aging and disease pathology.

**Figure 5.**
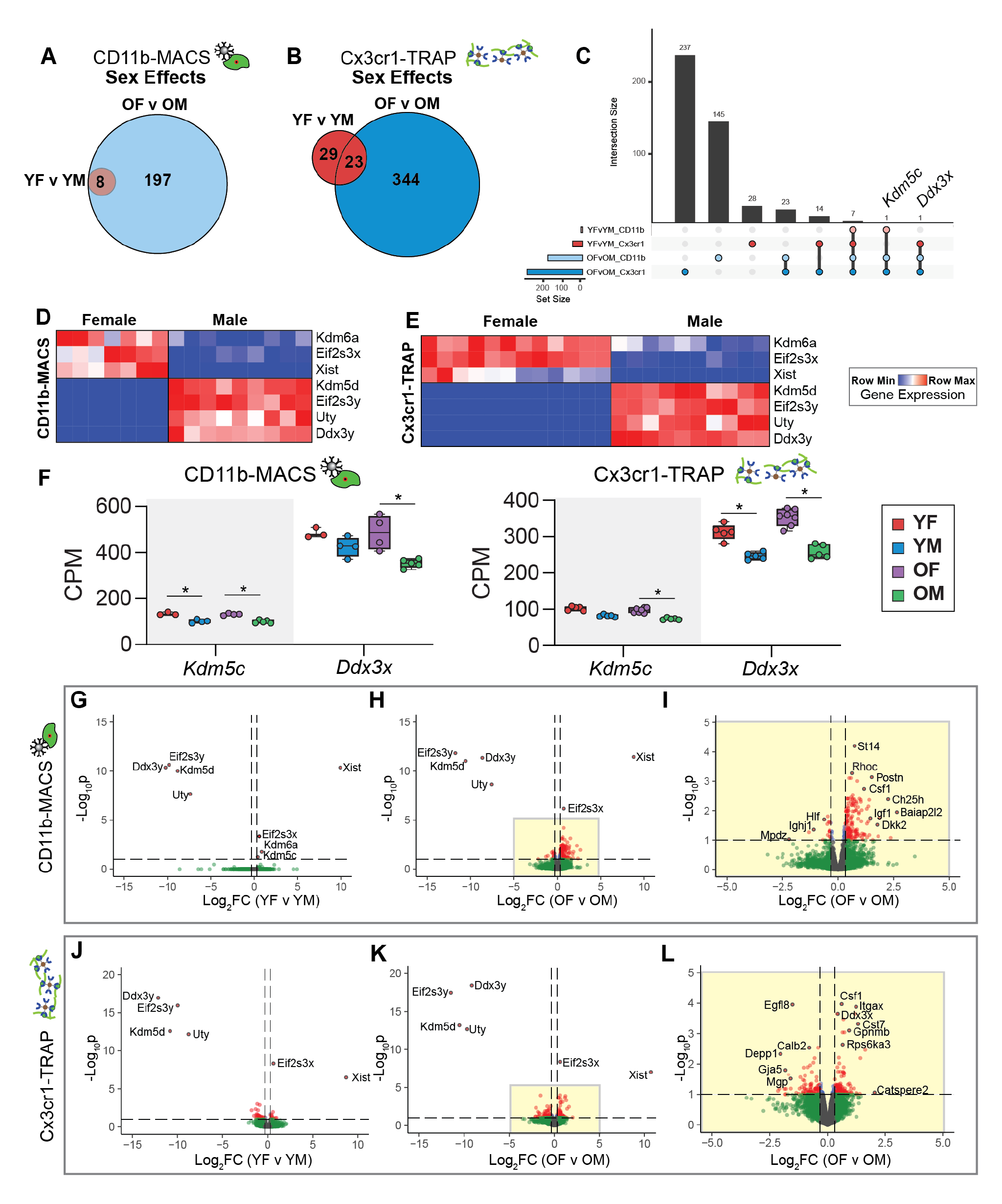
Analysis microglial transcriptomic (CD11b-MACS) and translatomic (Cx3cr1-TRAP) sex effects from young and old mouse hippocampus. **A)** Sets of differentially expressed genes (GLM QLF-test, FDR<0.1) showing sex effects in young (YF v YM) or old (OF v OM) within the CD11b-MACS dataset were compared by Venn diagram. **B)** Sets of differentially expressed genes (GLM QLF-test, FDR<0.1) showing sex effects in young (YF v YM) or old (OF v OM) within the Cx3cr1-TRAP dataset were compared by Venn diagram. **C)** Upset plot of differentially expressed genes by sex from both age groups and across methods (GLM QLF-test, FDR<0.1, |FC|>1.25). **D-E)** Heatmap of the gene expression of seven DEGs by sex that were identified in young and old mice with both **D)** CD11b-MACS and **E)** Cx3cr1-TRAP methodologies. **F)** Gene Expression (CPM) of *Kdm5c* and *Ddx3x* from CD11b-MACS and Cx3cr1-TRAP analyses (GLM QLF-test, |FC|>1.25, **\***FDR<0.1). **G-I)** Volcano plots of differentially expressed genes with sex (GLM QLF-test, FDR<0.1, |FC|>1.25) within CD11b-MACS for **(G)** young and **(H)** old mice. **I)** Inset of **Figure 5H** volcano plot showing differentially expressed genes between old females and old males within the CD11b-MACS analysis. **J-L)** Volcano plots of differentially expressed genes with sex (GLM QLF-test, FDR<0.1, |FC|>1.25) within Cx3cr1-TRAP for **(J)** young and **(K)** old mice. **L)** Inset of Figure 5K volcano plot showing differentially expressed genes between old females and old males within the CD11b-MACS analysis.

There were two genes (*Kdm5c*, *Ddx3x*) that were differentially expressed by sex in three of the four comparisons (**Figure 5C**). *Kdm5c* is an X-chromosome encoded histone lysine demethylase that acts on H3K4 (35). Defects in KDM5C activity result in X-linked intellectual disability in males (36) and can result in mild intellectual disabilities in female carriers (37). DEAD-box helicase 3 X-linked (*Ddx3x*) is an ATP-dependent RNA helicase and another cause of X-linked intellectual disability in both males and females (38). When we examined the expression levels of *Kdm5c* and *Ddx3x* across our analyses, we noted overall higher levels of expression within females, consistent with an escape from X-chromosome inactivation (**Figure 5F**).

Volcano plots of DEGs by sex showed similar distributions of DEGs by age timepoint across both methods (**Figure 5G-L; Supplemental Table 2-3**) (QLF test, FDR<0.1, |FC|>1.25). In particular, there were very few sex DEGs in the young group, with the identified DEGs primarily from the previously-mentioned sex chromosomally-encoded genes (**Figure 5G, J**). In the old age group (OF v OM), there were many more autosomally-encoded DEGs by sex, with markers of disease-associated microglia (21) (i.e., *Csf1, Itgax, Cst7*) being more highly expressed in old females than old males (as identified in at least one methodology) (**Figure 5H-I, K-L**). Since the sex-chromosomally-encoded genes had large magnitude fold changes between males and females, an inset of **Figures 5H** and **5K** is given as **Figures 5I** and **5L**, respectively. Of note, Colony Stimulating Factor 1 (*Csf1*) is a component of the SASP, providing further evidence that female microglia adopt a senescent phenotype more robustly than male microglia in the aged hippocampus.

A previous study analyzing sex differences in the aged mouse (24 mo) forebrain, identified 37 sex DEGs with several genes involved in the APOE-driven expression of CCL3 and CCL4, including *Spp1*, *Gpnmb*, *Lgals3*, *Clec7a*, *Apoe, Ccl3,* and *Ccl4 (22)*. When compared to the present study, we also identified *Spp1* and *Ctse* as sex DEGs in aged hippocampal microglia, in addition to 26 sex DEGs exclusive to the hippocampus (**Supplemental Figure 2**). By focusing on a specific brain region (hippocampus), we were able to identify more sexually divergent DEGs in aged microglia, as further elaborated below.

### Sexually divergent microglial gene expression in the aging mouse hippocampus

Twenty-three sex DEGs were identified in the old age group in both the transcriptomic and translatomic analyses (**Figure 5C**), and not in either of the young comparisons. A heatmap of the 23 sex DEGs revealed 19 genes that were more highly expressed in old females than old males, 1 gene that was more highly expressed in old males than old females, and 3 genes that were discordant in directionality between the CD11b-MACS and Cx3cr1-TRAP methods (**Figure 6A**). Included in the female-biased DEGs identified in aged microglia were three important cluster of differentiation (CD) cell surface markers that play important roles in microglial function, including CD11c (*Itgax*), CD22 (*Cd22*), and CD282 (*Tlr2*). CD11c^+^ microglia are a minority cell population in the developing brain that differentiate from CD11c^-^ microglia during early perinatal life after engulfment of apoptotic neurons, independent of conventional microglial activation (39). CD11c^+^ microglia are the only microglia that express osteopontin (*Spp1*), which was shown to be important for the phagocytic and pro-inflammatory functions of CD11c^+^ cells (39). *Itgax* and *Spp1* were also both identified as markers of the DAM phenotype (21) and are more highly expressed in old female microglia than their male counterparts in the present study.

**Figure 6.**
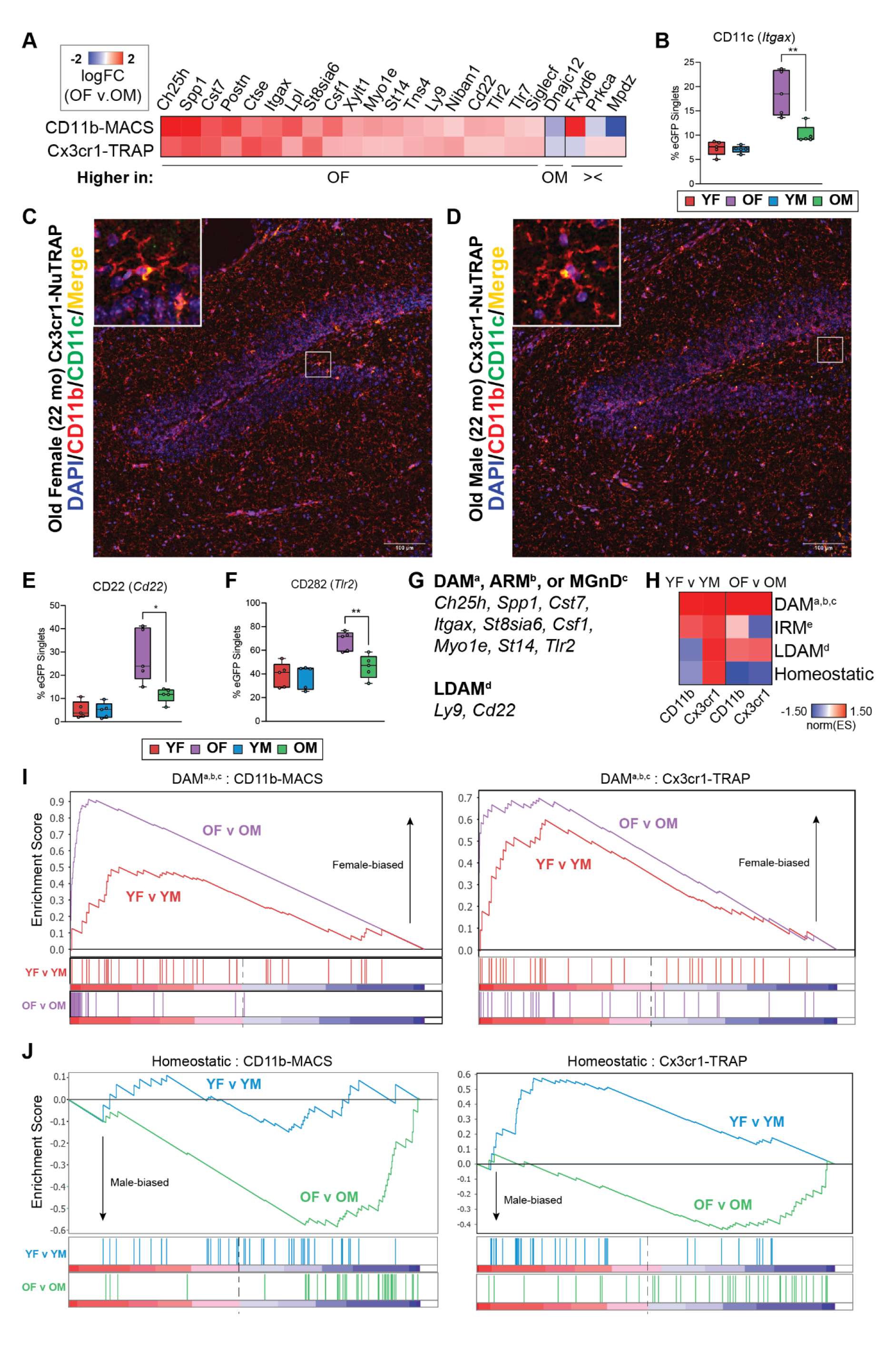
Sex differences in the transcriptome (CD11b-MACS) and translatome (Cx3cr1-TRAP) in old microglia. **A)** Heatmap of the logFC(OF/OM) from the 23 sex DEGs identified in the old age groups across both the transcriptome (CD11b-MACS) and translatome (Cx3cr1-TRAP). **B)** Flow cytometry confirmations of the percentage of CD11c (*Itgax*) (Two-way ANOVA, Sidak’s MTC, *p<0.05, **p<0.01, ***p<0.001). Boxplots represent the median +/- IQR. **C-D)** Representative confocal fluorescent microscopy images of sagittal brain sections captured in the hippocampus of old (22 mo) **C)** female and **D)** male mice show CD11c (pseudo-color green signal) in cells that co-expressed CD11b (red signal). **E-F)** Flow cytometry confirmations of the percentage of **E)** CD22 (*Cd22*), and **F)** CD282 (*Tlr2*) positive cells among the eGFP^+^ cells from Cx3cr1-NuTRAP hippocampus (Two-way ANOVA, Sidak’s MTC, *p<0.05, **p<0.01, ***p<0.001). Boxplots represent the median +/- IQR. **G)** List of genes with female-biased expression in old microglia that overlap with DAM^a^ (21), ARM^b^ (59), MGnD^c^ (60), and LDAM^d^ (42) markers. **H)** GSEA normalized enrichment scores comparing sex-biased enrichment of microglial phenotypic markers of DAM^a,b,c^ (21, 59, 60), IRM^e^ (59), LDAM^d^ (42), or homeostatic states in young (YF v YM) or old (OF v OM) groups. **I-J)** GSEA enrichment plots for **I)** DAM^a,b,c^ (21, 421, 59) and **J)** homeostatic marker genes.

To validate our transcriptomic findings at the protein level we performed flow cytometry on Cx3cr1-NuTRAP mouse hippocampus from young (5-6 mo) and old (22-25 mo) mice of both sexes (n=4-5/sex/age), using an independent cohort of mice from the transcriptomic and translatomic analyses. Consistent with our expression-level analyses, we observed a higher proportion of CD11c^+^ (**Figure 6B**). We also performed immunostaining of sagittal sections from Cx3cr1-NuTRAP brains from the original translatomic cohort and identified colocalized expression of CD11b and CD11c within microglia in the hippocampus of 22 mo male and female mice (**Figure 6C-D**). These representative images appear to show higher expression of CD11c in the old female microglia when compared to males.

CD22 was identified as a negative regulator of microglial phagocytosis in the aging brain (40) and was found to be upregulated in microglia in mouse models of AD (41). Further, disruption of CD22 signaling using a ligand-blocking antibody stereotactically injected into an aged mouse brain showed increased clearance of co-injected myelin debris or oligomeric amyloid beta (40). *Cd22* was also found to be upregulated in lipid-droplet accumulating microglia (LDAM), which are characterized by their accumulation of lipid-droplets and were found to have defective phagocytosis (42). On the other hand, CD282 (TLR2) is a cell surface pattern recognition receptor that was originally found to detect lipoteichoic acid and lipopeptide from gram-positive bacteria (43). In the brain, TLR2 detects damage-associated molecular patterns (DAMPs) produced during neuronal death and neurodegeneration (44). Of particular interest, TLR2 can be activated by fibrillar amyloid-beta to skew microglia to reactive phenotypes (45, 46). *Tlr2* was also found to be upregulated in the DAMs that have been associated with Aβ plaques.

We performed flow cytometry on Cx3cr1-NuTRAP mouse hippocampus from young (5-6 mo) and old (22-25 mo) mice of both sexes (n=4-5/sex/age), using the same cohort of mice as in **Figure 6B**. Consistent with our expression-level analyses, we observed a higher proportion of CD22^+^ (**Figure 6E**) and CD282^+^ (**Figure 6F**) cells among the eGFP^+^ microglia in old females when compared to old males, with no difference by sex within the young group.

We also noted that the expression of several genes that were more highly expressed in old female microglia was consistent with the DAM signature (*Ch25h*, *St8sia6i, Cst7, Lpl, Csf1, Myo1e, St14*) or LDAM signature (*Ly9*, *Cd22*) (**Figure 6G**). These results suggest that female microglia are adopting a disease-associated state with aging to a greater extent than male microglia, even in the absence of disease pathology. To assess the relative enrichment of microglial phenotypic markers in an unbiased way, we performed gene set enrichment analysis (GSEA) using DAM, interferon-response microglia (IRM), LDAM, and homeostatic marker genes compiled from multiple single-cell transcriptomic reports (**Supplemental Table 6**). We found a trend towards enrichment of DAM and LDAM phenotypic markers in old female microglia compared to male microglia, and a trend towards homeostatic markers in old male microglia when compared to female microglia (**Figure 6H**). Examination of the GSEA enrichment plots shows a greater level of enrichment of DAM markers in the OF v OM comparison than in the YF v YM comparisons by both methods (**Figure 6I**). Similarly, the GSEA enrichment plots showed a greater depletion of homeostatic microglial markers in the OF v OM comparison than in the YF v YM comparison by both methods (**Figure 6J**). These data further support the predicate that female hippocampal microglia skew towards reactive phenotypic states (i.e., DAM, LDAM) relative to males, with male microglia maintaining a more homeostatic transcriptomic in the aged brain.

### Pathway analysis of microglial transcriptomic and translatomic sex effects in the aging mouse hippocampus

To further interrogate the biological significance of the transcriptomic and translatomic sex effects in the aging mouse hippocampus, IPA was used to identify canonical pathways, diseases & biological functions, and upstream regulators that were differentially regulated by sex in hippocampal microglia at young and old age (**Figure 7A; Supplemental Table 7**). Within the canonical pathway analysis, we identified a sexually divergent induction of TREM1 signaling, NF- κB signaling, pathogen-induced cytokine signaling, and tumor microenvironment pathway. Of these pathways, there was no sex difference at young age, but the old females had greater induction of these pathways when compared to the old males. Triggering receptor expressed on myeloid cells 1 (TREM1) is a transmembrane receptor that can amplify inflammation through activation of the spleen tyrosine kinase (SYK) (47). Soluble TREM1 (sTREM1) is released in response to inflammation and can contribute to hippocampal neuronal damage and synaptic loss via the PI3K-AKT signaling pathway (48). NF-κB signaling can be upregulated by activation of PI3K-AKT signaling (49), suggesting that female-biased TREM1 activity may be upstream of the female-biased NF-κB upregulation in old microglia.

**Figure 7.**
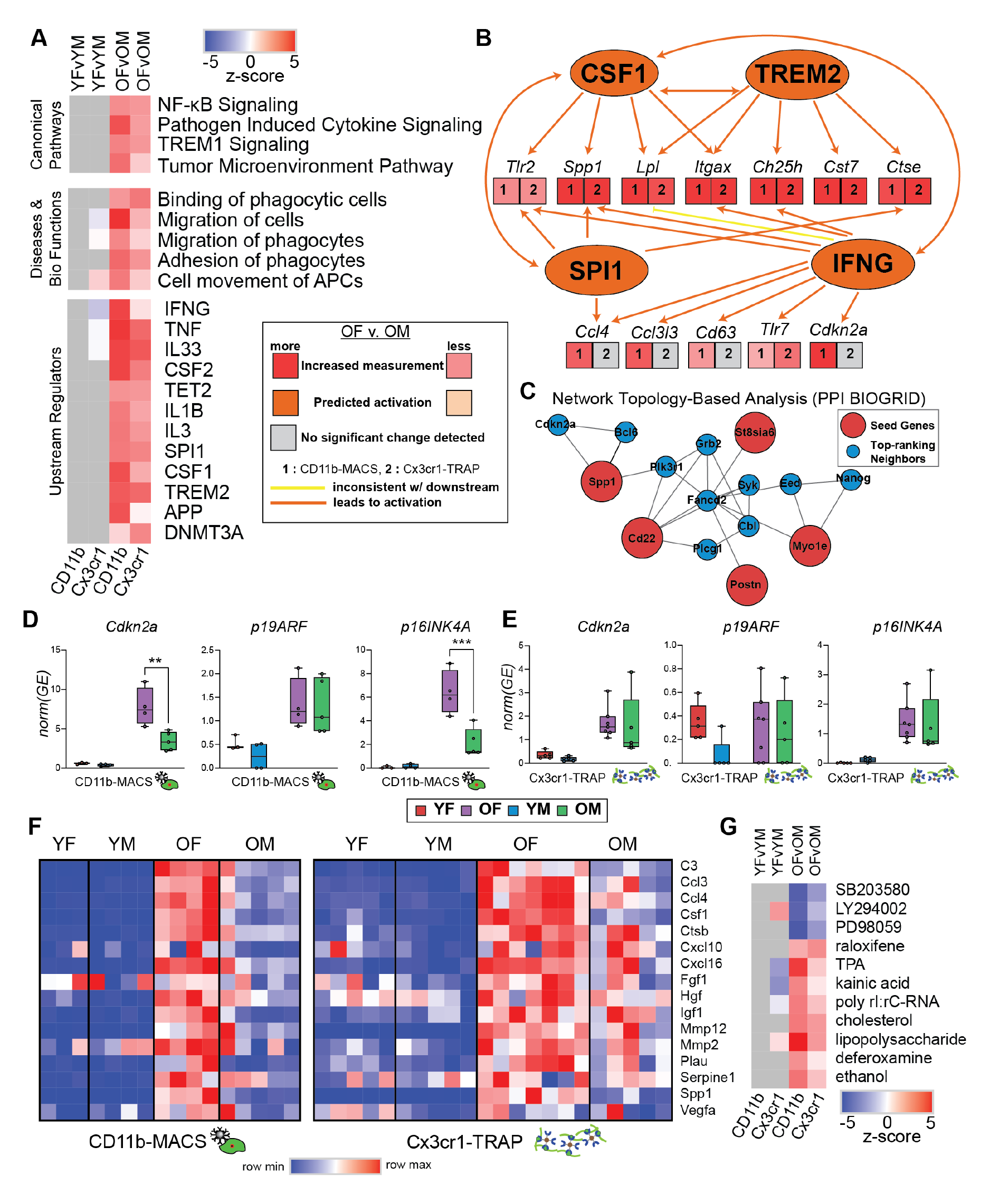
Pathway analysis of sex effects in the hippocampal microglial transcriptome and translatome. **A)** Heatmap of z-scores for IPA canonical pathways, diseases and bio functions, and upstream regulators analyses. **B)** IPA upstream regulator comparison to assess sex effects in old hippocampal microglia. **C)** Network topology-based analysis (PPI BIOGRID) of the 19 genes that were more highly expressed in old female microglia compared to old males (Figure 6A) in both the transcriptome and translatome. **D-E)** *Cdkn2a* transcript analysis for p19ARF and p16INK4A in the **D)** transcriptome and **E)** translatome of hippocampal microglia. **F)** Heatmap of SASP gene expression (CPM). **G)** IPA upstream regulator analysis to identify drug regulators of sex differences.

Within the diseases & biological functions pathways differentially regulated by sex, we identified several pathways involved in migratory and adhesion processes, suggesting sex differences in the motility of aged microglia. Examination of the differentially activated upstream regulators by sex identified many genes involved in pro-inflammatory response (i.e., IFNG, TNF, IL1B) in old female microglia. Interferon-gamma (IFNG) has been shown to prime microglia and cause a more responsive reaction to secondary stimulus, in addition to inducing microglial proliferation and altering hippocampal neuronal firing (50). Thus, higher induction of IFNG in old female microglia likely drives them more quickly towards senescence and subsequently results in higher susceptibility to AD pathology (i.e., amyloid). Tumor Necrosis Factors (TNFs), such as TNFα, are master regulators of pro-inflammatory cytokine release and have been shown to provide autocrine positive feedback to increase TNFα production in microglia (51). Higher activation of TNF pathways may drive this sex divergence of microglial inflammatory signaling through autocrine positive feedback loops, which may further be exacerbated with increasing age.

Another important upstream regulator that is more induced in old female microglia than old male microglia is SPI1, which encodes for the PU.1 transcription factor, a master regulator of microglial development and activation (52). SPI1 is also a genetic risk factor for AD (53) and has been linked to the proliferation of macrophages (52). Since SPI1 is a critical microglial transcription factor, even small changes in activity may lead to different susceptibilities to AD (52). Colony stimulating factor 1 (CSF1), another central regulator of microglial identity, showed higher activation in female microglia compared to old male microglia. CSF1 regulates the survival, proliferation, differentiation, and function of microglia and other macrophages (54). Since CSF1 induces proliferation (55), enhanced CSF1 signaling could lead to more exaggerated senescent phenotypes in female microglia and concomitant neuroinflammation. Female microglia also showed a larger activation of upstream regulator pathways associated with AD (i.e., APP, TREM2), suggesting a potential mechanism for female-biased disease progression in AD. Additionally, two epigenome modifier upstream regulators (TET2, DNMT3A) were shown to be more activated in old female microglia compared to old males, providing a potential epigenomic mechanism by which these sex differences in microglia may be perpetuated and maintained.

In addition to the solo activities of these upstream regulators that are more highly activated in old female microglia, several regulators provide positive feedback within a network (**Figure 7B**), such that the activation of one regulator further amplifies the effect of other regulators.

### Female-biased microglial senescence in the aging mouse hippocampus

We ran network topology-based analysis (PPI BIOGRID) on the 19 genes that were found to be more highly expressed in old female microglia when compared to old male microglia (**Figure 7C**). Among the top-ranking neighbors identified was *Cdkn2a*, the transcript encoding for p16INK4A, a well-established marker of senescence. (56). SYK was also identified as a top-ranking neighbor, and SYK inhibition was previously shown to be selectively toxic to senescent cells (57).

Since *Cdkn2a* encodes for two different transcripts with alternate start sites (p19ARF and p16INK4A), we performed transcript-specific analysis on the *Cdkn2a* transcriptome and translatome data. Within the CD11b-MACS transcriptomic data, we found that *Cdkn2a* was more highly expressed in old females compared to old males. This sex difference at old age was not observed in p19ARF expression, but instead was driven entirely by p16INK4A expression (**Figure 7D**). In the Cx3cr1-TRAP translatomic data, there was no sex difference observed in *Cdkn2a*, *p19ARF,* or *p16INK4A* expression (**Figure 7E**). In addition, there was a single old male that displayed higher expression of *Cdkn2a* and *p16INK4A* than any other samples in that group. ROUT outlier analysis (Q=5%) identified this as an outlier within the analysis. Since this sample was included in all other analyses, we did not remove it here but note that removal of this outlier results in a statistically significant difference between old females and old males in both *Cdkn2a* and *p16INK4A* expression. Therefore, we examined transcript-specific Cx3cr1-TRAP translatome data in an independent cohort of mice using RT-qPCR and confirmed that *Cdkn2a* expression was more highly expressed in the old females and was primarily driven by p16INK4A expression (**Supplemental Figure 3A**). We also used this same cohort to analyze telomere length as a possible marker of replicative senescence. Although telomere length was shorter in the positive (microglial) fraction compared to the mixed-cell input, there were no significant differences by age and/or sex identified (**Supplemental Figure 3B**). These results suggest that telomere shortening is not the source of the senescent phenotype observed in aged female microglia.

As another measure of senescence, we examined the expression of several previously-identified SASP factors (20) and noted an overall higher expression of SASP factors in old females when compared to all other groups (**Figure 7F**). Upstream analysis of sex DEGs showed several drug pathways that were differentially regulated by sex in the old group (**Figure 7G; Supplemental Table 7**). SB203580 (p38 MAPK inhibitor), LY294002 (PI3K inhibitor), and PD98059 (MEK inhibitor) were more highly induced in males. Thus, treatment with these drugs in females may abrogate the neuroinflammatory phenotypes observed. Greater induction of the lipopolysaccharide pathway in females (**Figure 7G; Supplemental Table 7**) might relate to the greater expression of toll-like receptors (*Tlr7, Tlr2*) in aged female microglia compared to males. Additionally, raloxifene (a selective estrogen receptor modulator) pathway was more induced in females than males with aging. This suggests that estrogen signaling may play a role in the sexually divergent response to aging in hippocampal microglia.

## Discussion

In the aging mouse brain, microglia adopt diverse phenotypic states, even in the absence of disease pathology (3). Some common characteristics of the transcriptomic changes identified in microglia with brain aging include the reduction in homeostatic marker gene expression and upregulation of genes associated with pro-inflammatory response (58). In addition, bulk transcriptomic and imaging studies in the mouse hippocampus have shown a greater upregulation of inflammatory mediators (i.e., complement, MHCI) with aging in females as compared to males (4, 5). A study in mouse forebrain identified sex divergences in the microglial translatomic response to aging that centered around APOE signaling (22). However, the contributions of microglia in the hippocampus to these sex effects could not be ascertained from this study. Additionally, recent single-cell transcriptomic studies have identified novel microglial phenotypic states (i.e., DAM, LDAM) observed with aging and disease pathology (21, 42, 59, 60). Bolstered with this new understanding of microglial diversity across age and disease, we sought to assess the relative contributions of microglial phenotypic states to the transcriptomic and translatomic sex effects observed in the aging mouse hippocampus. The data presented here provides insight into the mechanisms responsible for sex effects in microglial phenotypic responses to aging in the hippocampus, and how these basal differences may further be amplified in disease states, such as AD.

To ensure rigor and reproducibility, we used two validated methods for assessing microglial gene expression without inducing *ex vivo* activational confounds (7, 8) using acutely isolated hippocampal tissue from young and old mice of both sexes. Importantly, these studies recapitulated the microglial aging phenotypes observed in previous studies (3, 22), with upregulation of inflammatory signaling pathways, migratory functions, and age-related upstream regulators in the transcriptome and translatome of both sexes. The consistency of these findings further validate our methodologies for the identification of transcriptomic and translatomic changes in the aging mouse brain. These data are accessible through the searchable web interface https://neuroepigenomics.omrf.org/ as a resource for the field. The most important result of this study was the identification of sex divergences in the transcriptome and translatome of the aging mouse hippocampus. We found that female microglia more robustly adopt disease-associated (i.e., DAM, LDAM) phenotypic states and display a greater senescent signature than male microglia in the aged mouse hippocampus.

*Spp1* was found to be differentially expressed by sex in the old groups in our transcriptomic and translatomic analyses, as well as a previous study in the mouse forebrain (22). Of interest, *Spp1* is increased in microglia from mouse models of AD (21, 60) and is a marker of the DAM/ARM/MGnD phenotype. Recently, perivascular macrophage expression of *Spp1* was shown to be necessary for microglial engulfment of synapses and upregulation of phagocytic markers in the mouse hippocampus with amyloid pathology (61). *Spp1* was also identified as a component of the SASP, and is considered a marker of cellular senescence (20). This finding suggests that higher expression of *Spp1* in aged female microglia may lead to a greater susceptibility to neurodegeneration in AD.

Cathepsin E (*Ctse*) was also found to be differentially expressed by sex in the old groups in our transcriptomic and translatomic analyses, as well as a previous study in the mouse forebrain (22). *Ctse* encodes for CatE, a protease involved in the endosomal-lysosomal system, is exclusively expressed by reactive microglia, and increases in response to amyloid pathology (62). The transcription factor PU.1 (encoded by *Spi1*), which was identified as a female-biased upstream regulator in old microglia, enhances *Ctse* expression (63). CatE expression leads to the proteasomal degradation of autophagic degradation of inhibitor of κBα (IκBα) into NFκB, which allows NFκB to translocate to the nucleus to initiate pro-inflammatory signaling cascades (64). The female-biased *Ctse* expression in aged microglia may thus be contributing to the female-biased activation of NFκB signaling.

In addition to brain-region independent sex differences in the aged mice, we also identified hippocampal-specific sex differences in aged microglia, which included *Csf1*. Importantly, *Csf1* is a component of the SASP (20) and induces microglial proliferation (55). It stands to reason that female-biased upregulation of *Csf1* in aged microglia could lead to greater proliferation in female microglia and quicker transition to senescent phenotypes (6). *Csf1* has also been shown to be a p53 target that provides positive feedback to suppress apoptosis (65), further suggesting a role for *Csf1* in the female-biased senescent phenotype observed in the aged microglia analyzed in this study. Cystatin F (*Cst7*) was another hippocampal-specific sex DEG identified in the transcriptomic and translatomic analyses of the present study. *Cst7* is a marker of the DAM/ARM/MGnD phenotype (21, 59, 60) and involved in endolysosomal processing through inhibition of cysteine proteases (i.e., cathepsin L and C) (66). Interestingly, *Cst7* knockout had sexually divergent effects on microglia in the APP^NL-G-F^ mouse model of AD (67). Specifically, *Cst7* knockout in APP^NL-G-F^ mice led to the upregulation of endolysosomal genes in females and the downregulation of inflammatory genes in males. Female *Cst7*-deficient APP^NL-G-F^ mice also showed greater lysosomal and amyloid beta burden than their male counterparts. It may be the case that sex differences in other endolysosomal genes (i.e., *Ctse*) may lead to the sex differential response of microglia to *Cst7* deficiency in the context of amyloid pathology. These results highlight the importance of sex-specific analyses in elucidating aging and disease mechanisms, as different mechanisms may be operating as a function of sex.

Toll-like receptor 7 (*Tlr7*) was also found to be differentially expressed by sex in both the transcriptomic and translatomic analyses of old microglia. *Tlr7* is an endosomal immune sensor that detects single-stranded RNA and can elicit a type I interferon response to viral stimulus (68). *Tlr7* expression can be induced by NFκB activity (68) which, as previously mentioned, also has female-biased activity in aged microglia. In the kidney, small endogenous (non-viral) RNAs were able to activate *Tlr7* in the context of healthy aging (69). It is possible that the same mechanism is acting in aged microglia. *Tlr7* is an X-chromosomally encoded gene that variably escapes from X-chromosome inactivation. The higher levels of *Tlr7* in aged female microglia could be a result of escape from X-chromosome inactivation. Since *Tlr7* is expressed on the endosome, it is also possible that there are interactions with *Ctse* and *Cst7* to further exacerbate sex effects in aging microglia.

We showed that aged female microglia are enriched for DAM/ARM/MGnD markers (21, 59, 60) when compared to aged male microglia. We further showed female-biased activation of TREM2 in aged microglia. When TREM2 binds its ligand, it causes phosphorylation of immunoreceptor tyrosine-based activation (ITAM) receptor DAP12, which recruits Syk kinase to signal to downstream effectors (i.e., PI3K) which effect various essential cell functions (survival, proliferation, phagocytosis, motility) (70-72). TREM2 deficiency in mice prevents microglial clustering around amyloid beta plaques in the 5XFAD model (73) and prevented microglial transition to the Stage 2 DAM phenotype (21) and concomitant upregulation of *Cst7*, *Lpl*, *Csf1*, and *Itgax* (all identified as sex divergences in the present study). TREM2 signaling also interacts with CSF1R signaling through interaction with DAP12 and SYK signaling (71), which further supports the idea of multiple upstream regulators acting synergistically to promote and maintain the sexually divergent reactive phenotypes observed in the aging brain in health and disease. Recently, SYK signaling was shown to be required for the microglial phagocytic clearance of amyloid pathology in a mouse model of AD (74). Together, these data suggest that the emergence of the DAM phenotype is protective against destructive amyloid pathology. The female-biased overrepresentation of DAM markers may provide a cellular mechanism by which women with Alzheimer’s disease survive longer than men, but with worse pathology (75).

Recent AD clinical trials have shown that biological sex interacts with genomic risk factors to alter therapeutic efficacy, highlighting the need for precision medicine. For example, results of the Phase 3 Lecanemab trial revealed that females and APOE4 homozygotes did not show improved cognition in response to Lecanemab treatment, while males and APOE4 carriers showed significant improvement (76). Additionally, hormone replacement therapy was shown to interact with APOE genotype to yield different efficacies in AD risk prevention in post-menopausal women (77). Since APOE signals through TREM2 (60), the sex biases observed in TREM2 signaling pathways in aged microglia may be contributing to the differential response to AD treatment.

Although the data presented here robustly identifies a skew towards disease-associated and senescent phenotypes in aged female microglia, single-cell techniques will be needed to confirm sex differences in heterogeneity. In addition, validation of the senescent signature using traditional approaches (i.e., SA-β-gal staining) will need to be conducted. Mechanistic studies are needed to determine the function of individual genes in promoting the sexually divergent phenotypes described here.

Future studies will focus on the interactive effects of sex chromosomal (i.e., escape from X-chromosome activation) and sex hormonal (i.e., estrogen signaling) mechanisms in mediating sex effects in hippocampal microglial reactivity with aging and in models of Alzheimer’s disease. *Kdm6a* is a histone lysine demethylase that was found to be differentially regulated by sex across all comparisons in the present study and was shown to serve a protective effect in an AD mouse model (32). However, the role of *Kdm6a* and sex-specific histone modifications have not been explored in microglia with aging or in response to amyloid pathology. In addition, the sex differential regulation of DNA modifiers (DNMT3A, TET2) suggests that DNA methylation and hydroxymethylation may regulate the sex effects observed in this study. Finally, the identification of raloxifene (a selective estrogen receptor modulator) in the drug analysis suggests that estrogen signaling may also play a role in the observed sex effects. Interrogation of microglial-specific estrogen signaling will be needed to disentangle these effects.

## Conclusions

In conclusion, the data presented here identify novel genes that are differentially regulated by sex in the aged mouse hippocampus. The female-biased genes and pathways suggest that female microglia skew to disease-associated and senescent phenotypes, even in the absence of disease pathology. The pathways identified are highly relevant to AD and may be used to identify novel sex-informed, disease-modifying therapies. The transcriptomic and translatomic findings presented here are available in searchable format at https://neuroepigenomics.omrf.org/.

## Declarations

### Ethics approval and consent to participate

Not applicable

### Consent for publication

Not applicable

### Availability of data and materials

The datasets generated and/or analyzed during the current study are available in the NCBI Gene Expression Omnibus (GEO) repository, under accession number GSE233400. The transcriptomic and translatomic data can be accessed via a searchable web interface at https://neuroepigenomics.omrf.org/. Supporting data are provided as Supplemental Tables. All other data is available upon reasonable request from the corresponding author.

### Authors’ contributions

SRO designed the work; designed and performed experiments; acquired, analyzed, and interpreted data; and wrote the manuscript. KDP, JEJC, AWK, SK FAA, HLP, VAA, AK, CMK, AHM, and AJCE performed experiments; acquired, analyzed, and interpreted data; and revised the manuscript. WMF designed the work; designed experiments; analyzed and interpreted data; and substantively revised the manuscript.

## Acknowledgments

The authors would also like to thank the Clinical Genomics Center (OMRF), Imaging Core Facility (OMRF), Center for Biomedical Data Sciences (OMRF), and DMEI (OUHSC), and Flow Cytometry and Cell Sorting Core Facility (OMRF) for assistance and instrument usage.

## Authors’ information (optional)

Not applicable

---

**Supplemental Figure 1.**
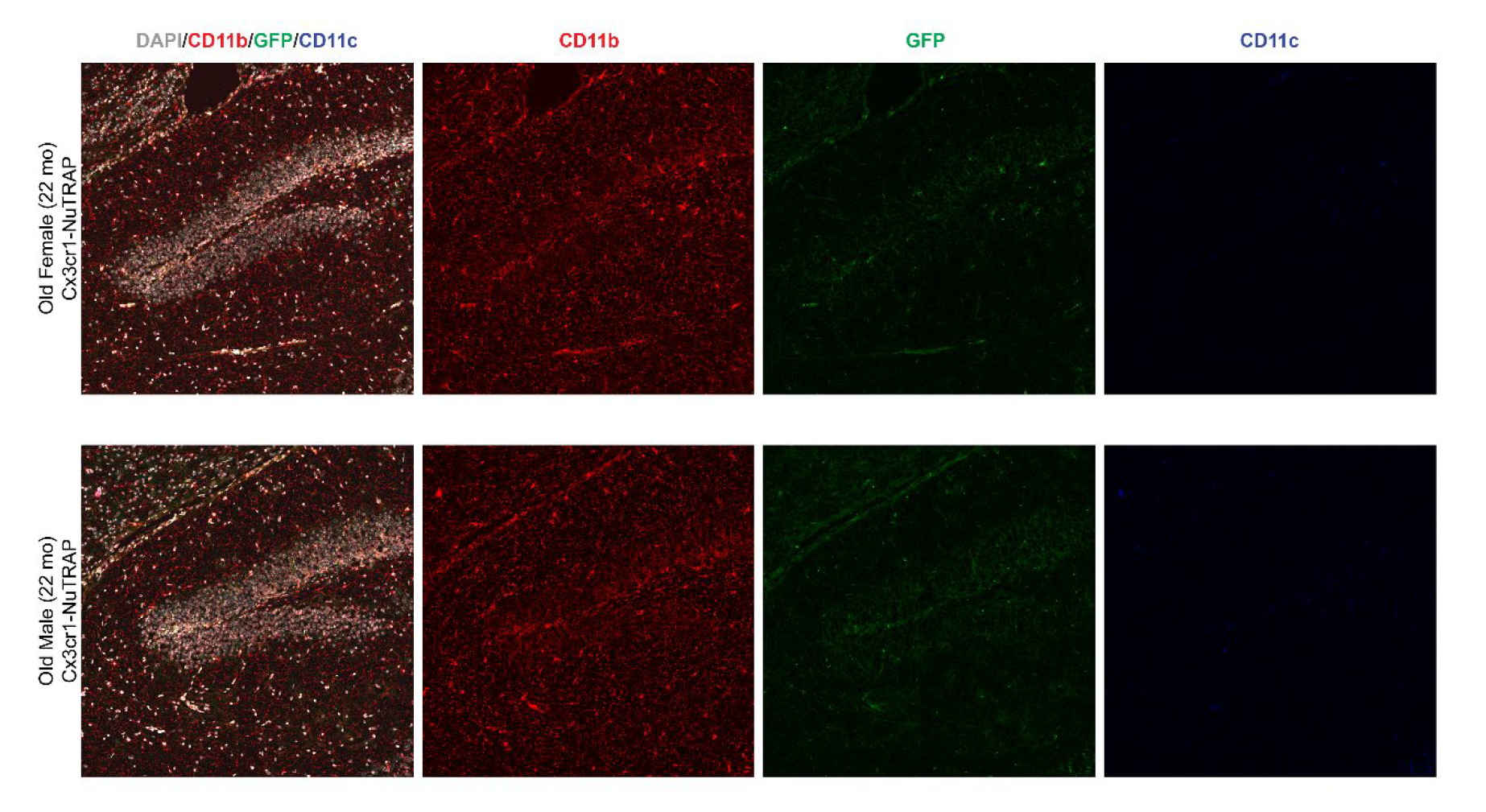
**Separated channels for IHC images displayed in Figure 6**. Cx3cr1-NuTRAP brains were processed for IHC analyses of frozen sections immunostained with antibodies against CD11b (red signal) and CD11c (blue signal). DAPI counterstaining of nuclei is indicated in grey. For the merged images presented in Figure 6, the eGFP channel was eliminated and the CD11c staining was pseudo-colored to green for better visualization of the colocalization of CD11b and CD11c, and DAPI was colored blue.

**Supplemental Figure 2.**
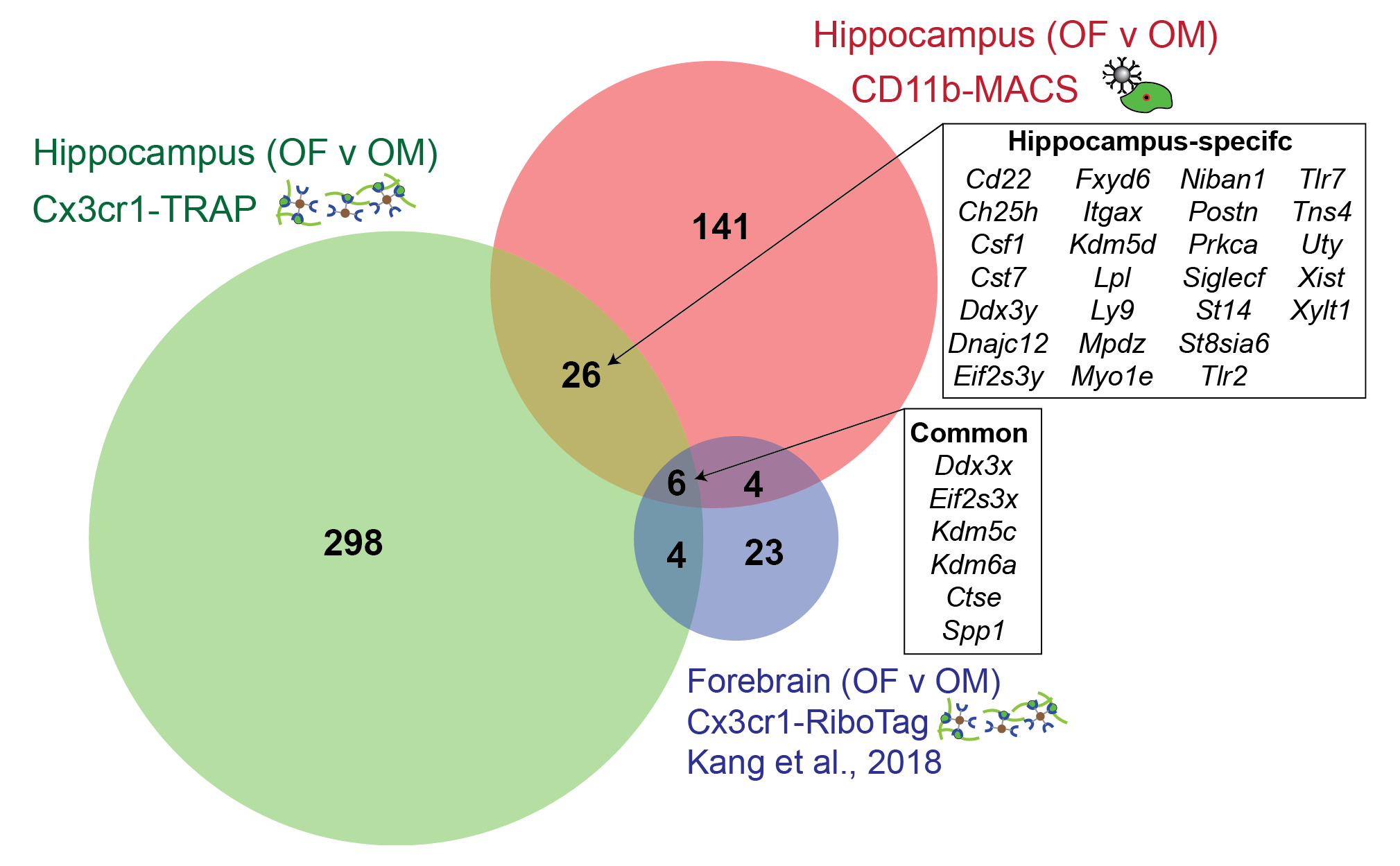
Comparison of sex effects in aged microglia (22-25 mo) from the present study (hippocampus) and a previously published study (Kang et al., 2018; forebrain) identifies hippocampus-specific sex effects.

**Supplemental Figure 3.**
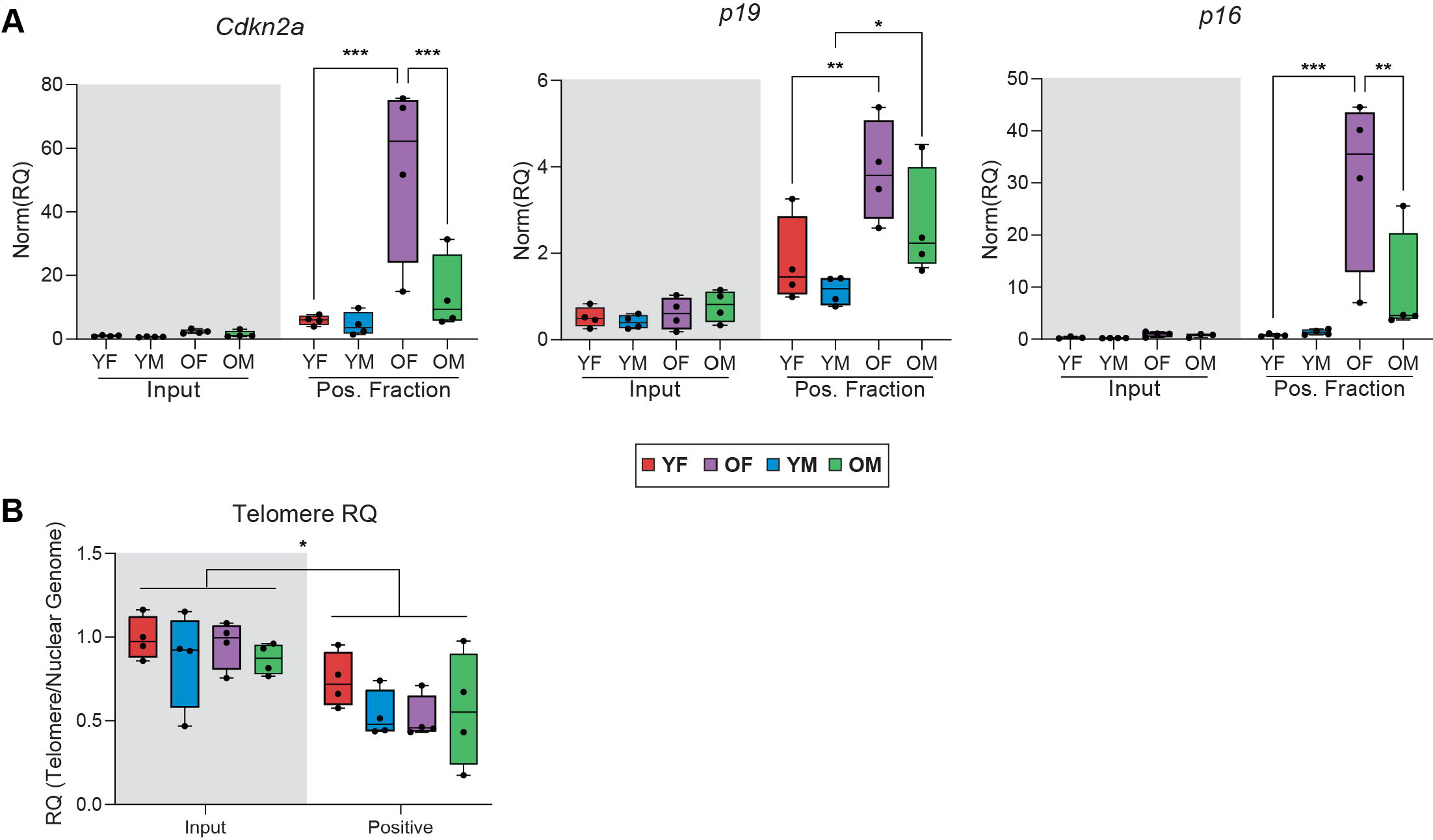
**Senescent marker measurement from the Cx3cr1-NuTRAP brain.** A separate cohort of male and female Cx3cr1-NuTRAP mice were aged to 8-13 mo and 23-26 mo (n=4/sex/age) for TRAP-RT-qPCR and INTACT telomere assays. **A)** RT-qPCR for *Cdkn2a* and its transcripts *p19* and *p16* (Two-way ANOVA, Tukey’s post-hoc, *p<0.05, **p<0.01, ***p<0.001). **B**) Relative telomere length as assessed by RT-qPCR (Two-way ANOVA, main effect of INTACT fraction (input v. positive), *p<0.05). Box plots represent median +/- IQR.

